# A two-locus system with strong epistasis underlies rapid parasite-mediated evolution of host resistance

**DOI:** 10.1101/2020.06.11.145391

**Authors:** Camille Ameline, Yann Bourgeois, Felix Vögtli, Eevi Savola, Jason Andras, Jan Engelstädter, Dieter Ebert

## Abstract

Parasites are a major evolutionary force, driving adaptive responses in host populations. Although the link between phenotypic response to parasite-mediated natural selection and the underlying genetic architecture often remains obscure, this link is crucial for understanding the evolution of resistance and predicting associated allele frequency changes in the population. To close this gap, we monitored the response to selection during epidemics of a virulent bacterial pathogen, *Pasteuria ramosa*, in a natural host population of *Daphnia magna*. Across two epidemics, we observed a strong increase in the proportion of resistant phenotypes as the epidemics progressed. Field and laboratory experiments confirmed that this increase in resistance was caused by selection from the local parasite. Using a genome wide association study (GWAS), we obtained a genetic model in which two genomic regions with dominance and epistasis control resistance polymorphism in the host. We verified this model by selfing host genotypes with different resistance phenotypes and scoring their F1 for segregation of resistance and associated genetic markers. Applying the model to the dynamics of the field population revealed moderate changes in allele frequencies at the two resistance loci relative to the profound changes observed at the phenotypic level. This apparent discrepancy is explained by strong epistasis and dominance at the two resistance loci, which reduces the effect of selection on alleles at both loci. Such epistatic effects with strong fitness consequences in host-parasite coevolution are believed to be crucial in the Red Queen model for the evolution of genetic recombination.

## Introduction

Darwinian evolution is a process in which the phenotypes that are best adapted to the current environment produce more offspring for the next generation. Genetic variants that code for these phenotypes are thus expected to increase in frequency in the population. Although this concept is fundamental in evolutionary biology, it remains difficult to connect the phenotype under selection with the underlying changes in the gene pool of natural populations (Ellegren and Sheldon 2008; Whitlock and Lotterhos 2015; Hoban et al. 2016). While single gene effects have been shown to explain the phenotype–genotype interplay in some naturally evolving populations (Daborn 2002; Cao et al. 2016; van’t Hof et al. 2016), the genetic architecture underlying a phenotype is often complex. In addition, the way the environment influences the expression of a trait, and genotype x environment interactions may further obscure the link between phenotype and genotype. It is, thus, often impossible to predict genetic changes in a population that result from selection on specific phenotypes. Among the most potent drivers of evolutionary change in host populations are parasites; parasite-mediated selection can raise the frequency of resistant phenotypes rapidly (Schmid-Hempel 2011, Morgan and Koskella 2017, Koskella 2018, Kurtz et al. 2016) and is thought to contribute to many biological phenomena, such as biodiversity (Laine 2009), speciation (Schlesinger et al. 2014) and the maintenance of sexual recombination in the host (Lively 2010, Gibson et al. 2018).

To link patterns produced by parasite-mediated selection with evolutionary theory, we need to know the genetic architecture that underlies resistance; this includes the number of loci, their relative contribution to the phenotype, and the interaction between loci (epistasis) and alleles (dominance). In this way, we can predict the outcome of selection, test theoretical models, and understand epidemiological dynamics (Hamilton 1980; Galvani 2003; Schmid-Hempel 2011). In a few cases, resistance to parasites has been found to be determined by single loci with strong effects, e.g. in plants (Gómez-Gómez et al. 1999; Li and Cowling 2003; Li et al. 2017), invertebrates (Juneja et al. 2015; Xiao et al. 2017), and vertebrates (Samson et al. 1996). However, the genetic architecture is often obscured by intrinsic complexity and confounding factors that may interact with the phenotype. Resistance might be determined by multiple loci with qualitative or quantitative effects, present distinct dominance patterns, and display interactions with other genes or the environment. Indeed, multi-locus genetic architecture of resistance is thought to be more common than single-locus, as it can create more diversity (Sasaki 2000; Tellier and Brown 2007; Wilfert and Schmid-Hempel 2008). Multi-locus architecture was described in *Drosophila melanogaster*, for example, where resistance was found to be determined mostly by a few large-effect loci (Bangham et al. 2008; Magwire et al. 2012) and some additional small-effect loci (Cogni et al. 2016; Magalhães and Sucena 2016). Quantitative resistance has also been found in crops where it can be used as a pathogen control strategy (Pilet-Nayel et al. 2017). In the water flea *Daphnia magna*, resistance has been found to be quantitative to a microsporidian parasite, but qualitative to a bacterial pathogen (Routtu and Ebert 2015). Although resistance tends to be dominant (Hooker and Saxena 1971, Carton et al. 2005), resistant alleles have been found to be both dominant and recessive in plants (Gómez-Gómez et al. 1999; Li and Cowling 2003; Li et al. 2017) and invertebrates (Luijckx et al. 2012; Juneja et al. 2015; Xiao et al. 2017). Epistasis between resistance loci has also been found in diverse plants and animals (Kover and Caicedo 2001; Wilfert and Schmid-Hempel 2008; Jones et al. 2014; González et al. 2015; Metzger et al. 2016), emphasizing its crucial role in the evolution of resistance. The link between genetic architecture and natural selection for resistance remains weak, however, mainly limited to the theoretical extrapolation of results from laboratory experiments.

Dominance and epistasis describe the non-additive interaction among alleles of the same or different loci, respectively, making them crucial for the evolutionary response to selection. Epistasis among resistance genes is thought to be pervasive, as it could contribute to the maintenance of genetic diversity by reducing fixation rates at individual loci (Tellier and Brown 2007). In the Red Queen model for the evolution of sex, thus, epistasis among resistance loci helps maintain genetic diversity and recombination in the host (Hamilton 1980; Hamilton et al. 1990; Howard and Lively 1998; Salathé et al. 2008; Engelstädter and Bonhoeffer 2009; Kouyos et al. 2009). The importance of genetic architecture for understanding the evolution of resistance stands in stark contrast to the limited amount of available data on natural populations (Alves et al. 2019). In this study, we investigate the evolution of resistance in a natural population of the planktonic crustacean *Daphnia magna*, as it experiences epidemics of the bacterial pathogen *Pasteuria ramosa*. We link parasite-mediated selection to its associated allele frequency change by resolving the underlying genetic architecture of resistance.

In recent years, water fleas of the genus *Daphnia* (Crustacea, Cladocera) and their microparasites have become one of the best understood systems for studying the evolution and ecology of host-parasite interactions (Ebert 2005, Vale et al. 2011, Izhar and Ben-Ami 2015, González-Tortuero et al. 2016, Strauss et al. 2017, Turko et al. 2018, Shocket et al. 2018, Rogalski and Duffy 2020). Parasite selection in natural *Daphnia* populations has been shown to alter the phenotypic distribution of resistance (Little and Ebert 1999; Decaestecker et al. 2007; Duffy and Sivars-Becker 2007; Duncan and Little 2007), and genetic research has identified loci involved in host resistance (Bento et al. submitted; Luijckx et al. 2012; Luijckx et al. 2013; Routtu and Ebert 2015; Metzger et al. 2016; Bento et al. 2017); however, because these studies largely involved crosses among populations, the results may not reflect genetic variation within populations. Genetic changes in natural host populations have been observed but not directly linked to parasite resistance loci (Mitchell et al. 2004; Duncan and Little 2007). Understanding the link between parasite-mediated selection on host resistance and the underlying genetic architecture would enable us to determine the tempo and mode of evolution in natural populations and to link observed phenotypic changes to frequency changes of alleles under selection. This study provides such a phenotype-genotype link. We quantified the change in frequency of resistance phenotypes over time in a natural *D. magna* population and, through experiments, showed that the locally dominant, virulent parasite genotype of *P. ramosa* played a major role in the observed phenotypic changes. A genome-wide association study (GWAS) and genetic crosses revealed the underlying genetic architecture of resistance in our study population and provided a genetic model for inheritance of resistance. Using this model, we were able to predict the changes in allele frequencies in our study population at two resistance loci during parasite-mediated selection. Dominance and epistasis caused only a comparatively weak change in allele frequency, despite strong selection on phenotypes. In summary, our study describes strong parasite-mediated natural selection on host resistance, and shows how this selection alters the genetic structure of the host population.

## Results

### Parasite-mediated selection explains phenotypic dynamics

In the Aegelsee *Daphnia magna* population, which we monitored from fall 2010 to fall 2015, *D. magna* diapauses during winter as resting eggs, while the active season spans from early April to early October. Each summer, we observed a *Pasteuria ramosa* epidemic that typically started in early May, about a month after *Daphnia* emerged from diapause, and lasted throughout the summer (Fig. 1A) with peak prevalence of 70% to 95%. *P. ramosa* infection in the host is characterized by gigantism, a reddish-brownish opaque coloration, and castration, i.e. an empty brood pouch. *P. ramosa* is a virulent parasite, stripping the host of 80% to 90% of its residual reproductive success and killing it after six to ten weeks, at which point it releases millions of long-lasting spores into the environment (Ebert et al. 1996; Ebert et al. 2016). In 2014 and 2015, we determined resistotype (resistance phenotype) frequencies of *D. magna* in this population relative to four *P. ramosa* isolates: C1, C19, P15 and P20. P20 had been isolated from our study population in May 2011; isolates C1, C19 and P15 had previously been established in the laboratory from other *D. magna* European populations.

**Figure 1.**
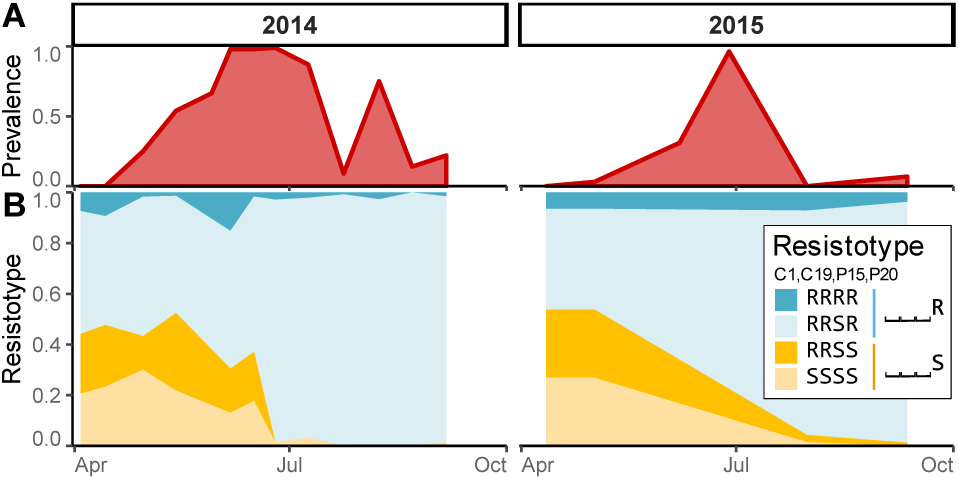
Dynamics observed in the Aegelsee *Daphnia magna* population. **A**: *Pasteuria ramosa* prevalence across strong summer epidemics. **B**: Resistotype (resistance phenotype) frequencies across time for 60-100 *Daphnia* clones from each sampling date in two-to three-week intervals from early April to early October 2014 and 2015. Resistotypes = resistance to *P. ramosa* C1, C19, P15 and P20, consecutively).

In 2011, we sampled a subset of infected animals (n = 113) to characterize *P. ramosa* diversity among infected hosts throughout the active season and found that the P20 genotype represented about 50% of the parasite diversity among infected hosts when the epidemics began. This proportion decreased to zero in mid-June, as other *P. ramosa* genotypes took over (Supplementary *Pasteuria* Fig. S1).

*D. magna* sampled from the field were cloned, and their resistotypes were scored. *D. magna* produces asexual clonal eggs which are used in the laboratory to produce clonal lines, a.k.a. genotypes. Individuals castrated by the parasite received an antibiotic treatment to allow clonal reproduction. Resistance to the bacteria is indicated when parasite spores are unable to attach to the gut wall of the host (Duneau et al. 2011; Luijckx et al. 2011). We thus defined host clone resistotype according to the ability of given isolate bacterial spores to attach to the host gut wall or not. The host’s overall resistotype is its combined resistotypes for the four individual *P. ramosa* isolates in the following order: C1, C19, P15 and P20, e.g. a clone susceptible to all four isolates will have a SSSS resistotype. When one isolate was not considered, we used the placeholder “⎵” for that resistotype: e.g. “RR⎵R resistotype”. Overall, we found found three predominant resistotypes: RRSR, RRSS and SSSS, which together accounted for 95.1±1.0% of all tested animals over the active season in 2014 (n = 995) and 2015 (n = 260). RRRR represented a much smaller proportion of the resistotypes (4.9±1.0%) (Fig. 1B). Excluding the data for *P. ramosa* isolate P15, for which over 95% of the hosts were susceptible, the study population was mainly composed of three resistotypes: RR⎵R, RR⎵S and SS⎵S. A few other resistotypes that were absent in the 2014 and 2015 samples were observed in other samples. Notably, the SR⎵S resistotype was found in 0.3% of hatched animals from *D. magna* resting eggs sampled during the winter 2014 diapause. The SR⎵R resistotype has never been found in the field samples, but was found in the selfed offspring of the rare resistotype SR⎵S. Resistotypes RS⎵⎵ and SS⎵R were not observed in this population.

The temporal dynamics revealed an increase in animals resistant to P20 (RRSR and RRRR, in short: RR⎵R, or ⎵⎵⎵R) soon after the onset of the epidemics, while animals susceptible to P20 (RRSS and SSSS, or ⎵⎵⎵S) declined accordingly (Fig. 1B) in both study years. Resistance to C1, C19 and P15 did not seem to play a strong role in the selection process during the epidemics. In the following results, we tested the hypothesis that selection by *P. ramosa* isolate P20 is the main driver of natural resistotype dynamics in our study population.

First, we tested the impact of the parasite on the different resistotypes to associate disease phenotype with resistotype. To do this, we obtained a sample of the spring cohort of the *D. magna* population by hatching resting eggs collected in February 2014. These animals represented, in total, 70 clones of the four most common resistotypes (RRSR, RRSS, SSSS, RRRR), each replicated five times. We exposed these clonal offspring to a mixture of *P. ramosa* spores that represented the diversity of the parasite population during the early phase of the epidemic. We then tested the hosts for infection (looking for visible signs) and its effect on fecundity (counting the number of produced clutches). Sixteen animals died before we could test their infection status, resulting in a total sample size of n = 334. Individuals with resistotypes RRSS and SSSS (susceptible to P20) were infected far more frequently than RRSR and RRRR (resistant to P20) individuals (Fig. 2A; Null deviance = 461.3 on 333 df, Residual deviance = 358.4 on 329 df, p < 0.001). The analysis also compared P20-susceptible and P20-resistant resistotypes, confirming the high susceptibility of the P20-susceptible animals (Fig. 2A; Null deviance = 461.3 on 333 df, Residual deviance = 360.7 on 331 df, p < 0.001). Infected P20-susceptible individuals produced on average about one less clutch before parasitic castration (n = 136, 1.83 ± 0.07 clutches) than did infected P20-resistant individuals (n = 19, 2.53 ± 0.3 clutches) (Fig. 2B; Null deviance = 76.9 on 154 df, Residual deviance = 74.0 on 152 df, p = 0.023). Accordingly, the average time period until visible infection was shorter in P20-susceptible clones (15.7 ± 0.2 days) than in P20-resistant clones (19.4 ± 1 days) (Fig. 2C; Null deviance = 94.4 on 154 df, Residual deviance = 85.1 on 152 df, p = 0.0018). These results clearly support the hypothesis that early season *P. ramosa* strains from the field select on the P20-resistotype.

**Figure 2.**
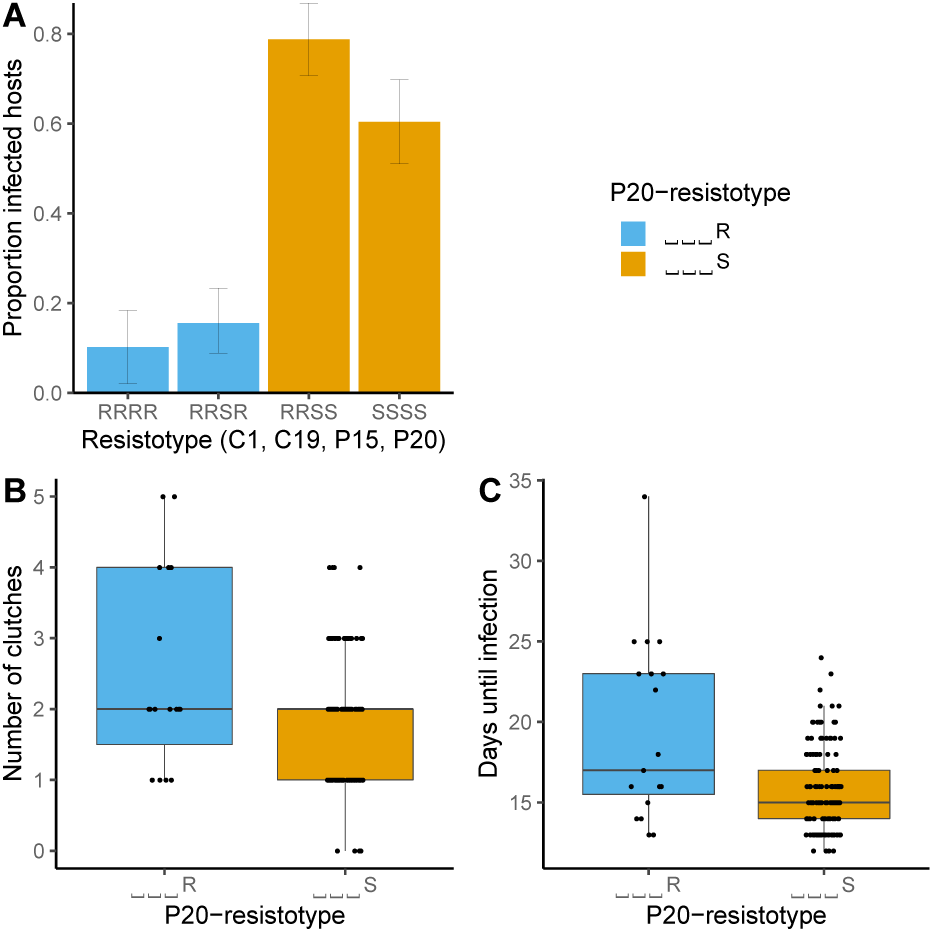
Experimental infections of *Daphnia magna* with different resistotypes (resistance phenotype). Resistotypes RRSR, RRSS, SSSS (n = 20 clones for each) and RRRR (n = 10 clones) were infected with parasite spores from the early phase of the epidemic. Five repeats were performed for each clone (total n = 334). Controls (n = 210) remained uninfected and are not shown here. **A**: Proportion of infected *Daphnia* among the four resistotypes. **B**: Number of clutches produced before parasitic castration in the infected P20-resistant (⎵⎵⎵R) and susceptible (⎵⎵⎵S) animals (n = 115). **C**: Time before visible infection in P20-resistant and P20-susceptible individuals (n = 115).

In the following year, we looked at the relationship of disease phenotype and P20-resistotype in the field by measuring the parasite’s impact on the different resistotypes. We collected animals in the field during the early half of the *P. ramosa* epidemic and raised them individually in the laboratory, recording their disease symptoms. We then cured infected animals with antibiotics, allowed them to produce clonal offspring, and determined their resistotype. Our analysis revealed higher infection rates (size corrected) for P20-susceptible individuals than for P20-resistant individuals in these natural conditions (Fig. 3A; Fitted model: glm (Infected (1/0) ∼ P20-resistotype + Body_size + Sampling_date, family = quasibinomial(), n = 331; Null deviance = 415.1 on 330 df, Residual deviance = 209.1 on 327 df, p = 0.025). Field-caught P20-susceptible individuals also produced, on average, fewer offspring before castration than P20-resistant ones (Fig. 3B; Fitted model: glm.nb (Fecundity ∼ P20-resistotype + Body_size * Sampling_date), n = 224; Null deviance = 127.9 on 223 df, Residual deviance = 92.9 on 219 df, p = 0.014). The data differences are partially due to the difference in parasite prevalence on the two sampling dates (31% on 7 June and 96% on 28 June 2015). These results also support the hypothesis that early season *P. ramosa* strains from the field select on the P20-resistotype.

**Figure 3.**
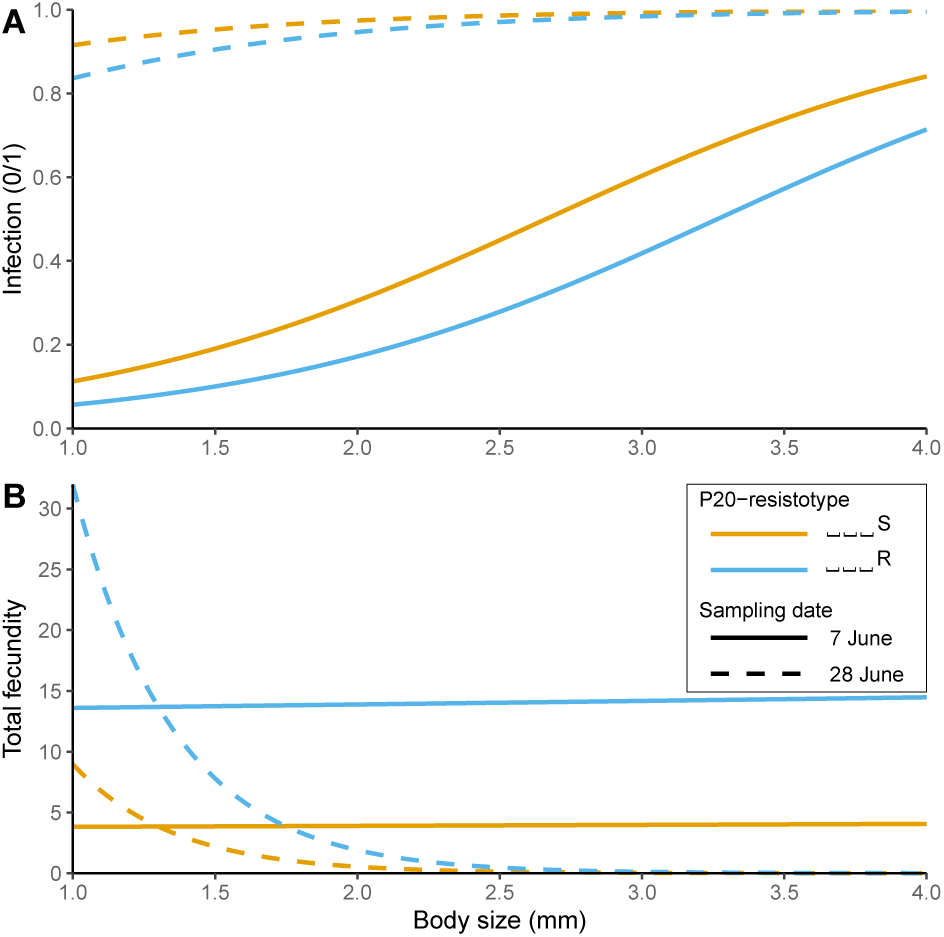
Fitted models of infection phenotypes in field-collected *Daphnia magna* relative to their body size at capture (x-axis) and their resistance to P20 for two sampling dates in June 2015. **A**: P20-susceptible (orange) animals have a higher likelihood to be infected than P20-resistant (blue) ones for any body size. **B**: Infected P20-susceptible animals have a lower total fecundity than P20-resistant ones for any body size. Differences between the data are partially due to the difference in parasite prevalence on the two sampling dates (31% on 7 June and 96% on 28 June).

### Linking resistance phenotypes to genotypes

After excluding P15, which has very low variability because most animals are P15-susceptible, the study population was composed mainly of three resistotypes: RR⎵R, RR⎵S and SS⎵S. Very rarely did we find the SR⎵S resistotype, while RS⎵⎵ and SS⎵R were never observed in this population. A supergene for resistance to C1 and C19 has been described in *D. magna* using QTL (Routtu and Ebert 2015; Bento et al. 2017), and the genetic architecture of resistance at this so-called ABC-cluster, or *P. ramosa* resistance (PR) locus, has been further resolved using genetic crosses among host genotypes (Metzger et al. 2016). According to this genetic model, an SS⎵⎵ resistotype (susceptible to C1 and C19) has an “aabbcc” genotype (lower case letters indicate recessive alleles), while RS⎵⎵ individuals (resistant to C1 and susceptible to C19) are “A---cc” (upper case letters indicate dominant alleles and a dash “-” indicates alleles that do not influence the phenotype); SR⎵⎵ individuals are “aaB-cc”, and RR⎵⎵ individuals are “C-”. In other words, allele A epistatically nullifies variation at the B-locus, and allele C nullifies variation at the A- and B-loci (Metzger et al. 2016; Bento et al. 2017). See also Supplementary Model Fig. S2.

Considering this genetic model, we assume that the recessive allele at the A-locus is fixed in our study population (“aa” genotype) and that the dominant allele at the B-locus is very rare, as we never observed RS⎵⎵ individuals and only found SR⎵⎵ in very low proportions. In our population, the SS⎵⎵ /RR⎵⎵ polymorphism can be best described by the C-locus polymorphism, i.e. genotypes “aabbcc” and “aabbC-”, respectively, with C being the dominant allele for resistance. Given this, we assume, in the following sections, that variation at the C-locus underlies the resistance polymorphism for C1 and C19.

### Genomic regions of resistance to the parasite

We sequenced the genomes of 16, 10 and 11 clones with resistotypes RR⎵R, RR⎵S and SS⎵S, respectively and conducted a GWAS comparing five pairs of these resistotypes to identify candidates for resistance to C1, C19 and P20: (i) SS⎵⎵ vs. RR⎵⎵, (ii) SS⎵S vs. RR⎵S, (iii) ⎵⎵⎵S vs. ⎵⎵⎵R, (iv) RR⎵S vs. RR⎵R and (v) SS⎵S vs. RR⎵R. Comparison (i) between SS⎵⎵ and RR⎵⎵ (variation at C1- and C19-resistotype) revealed a strong signal on linkage group (LG) 3 (Fig. 4A). This region encompasses the super gene described earlier by Routtu and Ebert (2015) and Bento et al. (2017), the so-called ABC-cluster, or *P. ramosa* resistance (PR) locus. Comparison (iii) between ⎵⎵⎵R and ⎵⎵⎵S (variation at P20-resistotype) revealed a strong signal on LG 5 (Fig. 4B), hereafter called the E-locus. In the present host-parasite system, the D-locus determines resistance to P15 and is not considered here (Bento et al. submitted). The E-locus region has not yet been associated with resistance, and no *P. ramosa* resistance gene has been described on the same linkage group in *D. magna*. Finally, comparison (v) between RR⎵R and SS⎵S (variation at C1-, C19- and P20-resistotype) indicated a strong signal at both the ABC-cluster and the E-locus (Fig. 4C). Comparisons (ii) and (iv) were consistent with this pattern (Supplementary GWAS Fig. S3). The genomic regions associated with resistotypes in our GWAS were not sharp peaks, but rather table-like blocks of associated SNPs (Fig. 4). This structure was expected for the C-locus, which is a known supergene - a large block of genome space with apparently little or no recombination that contains many genes (Bento et al. 2017). Fig. 4 indicates that the same may be the case for the E-locus, where the block of associated SNPs makes up nearly half of the linkage group. A few single SNPs also showed significant association in all the comparisons (Supplementary GWAS Fig. S3), but because of the strength of the observed pattern and because we expected a large region to be associated with resistance, we do not consider these single SNPs.

**Figure 4.**
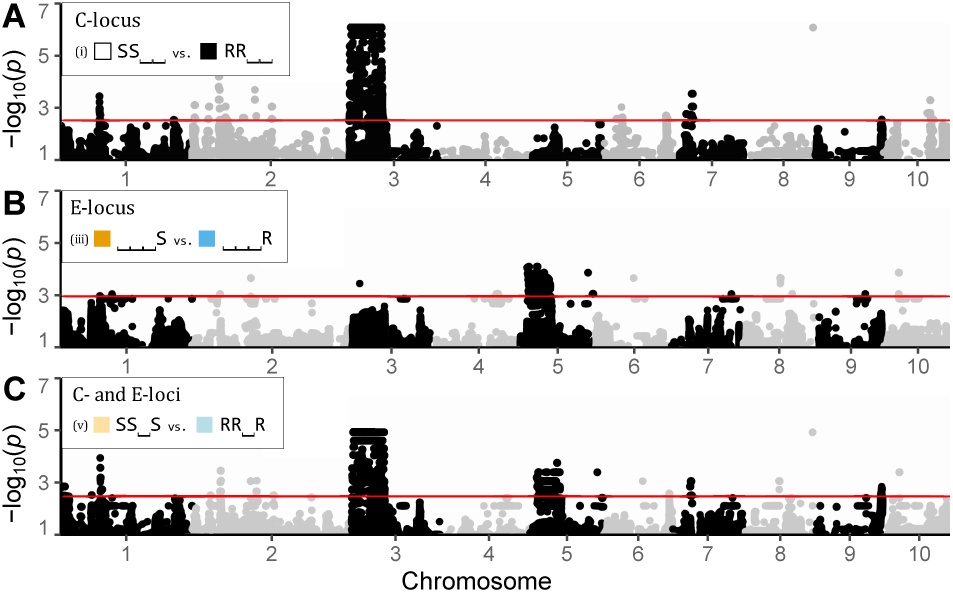
GWAS analysis comparing the most common resistance phenotypes (resistotypes) in the Aegelsee *Daphnia magna* population. Manhattan plots of relationships between different resistotype groups (showing only SNPs with *P*_*corrected*_ < 0.01). Resistotypes = resistance (R) or susceptibility (S) to C1, C19, P15 and P20 in that order. **A**: Comparison (i) SS⎵⎵ vs. RR⎵⎵ (variation at C1- and C19-resistotypes) revealed a strong signal on linkage group (LG) 3, corresponding to the C-locus. **B**: Comparison (iii), ⎵⎵⎵S vs. ⎵⎵⎵R (variation at P20-resistotype) revealed a strong signal on LG 5, corresponding to the E-locus. **C**: Comparison (v), SS⎵S vs. RR⎵R (variation at C1- and C19-, and P20-resistotypes), revealed a strong signal on both regions. The x-axis represents SNP data on the 2.4 *D. magna* reference genome and the genetic map (Routtu et al. 2014), showing only SNPs, not physical distance on the genome. The horizontal red line is the significance threshold, the maximum corrected p-value for which the q-value is inferior to 5% (see Methods section). Comparisons (ii) and (iv) are consistent with these results and are presented in Supplementary GWAS Fig. S3.

The E-locus region encompassed 22 scaffolds and one contig on version 2.4 of the *D. magna* reference genome, with a cumulative length of more than 3 Mb (3101076 bp) (Supplementary GWAS Table S1). We found 485 genes on all associated scaffolds. The strongest signals of association were found on scaffolds 2167 and 2560, which harboured 82 genes. Some of these genes were similar to genes identified in a previous study of the ABC-cluster on LG 3 (Bento et al. 2017), with a glucosyltransferase found on scaffold 2167. Three other sugar transferases (galactosyltransferases) were identified, two of them on scaffold 2560 (Supplementary GWAS Table S2).

### Genetic model of resistance inheritance

Mean allele frequencies at associated SNPs showed that SS⎵⎵ individuals (susceptible to C1 and C19) display a single allele at the C-locus, while RR⎵⎵ individuals display two distinct alleles at the C-locus. In other words, SS⎵⎵ individuals are homozygous at the C-locus, while RR⎵⎵ individuals comprise homo- and heterozygotes. At the E-locus, ⎵⎵⎵R individuals (resistant to P20) are homozygous, while ⎵⎵⎵S individuals (susceptible to P20) comprise homo- and heterozygous individuals (Supplementary GWAS Fig. S3). These results indicate that resistance to C1 and C19 is governed by a dominant allele (“C-” genotype), as shown before (Metzger et al. 2016). In contrast, resistance to P20 is determined by a recessive allele (“ee” genotype). Screening individual genomes revealed that some SS⎵S individuals (susceptible to C1 and C19, and P20) present the “ee” genotype at the E-locus (underlying P20-resistotype), although this genotype should confer resistance to P20. This was not observed in RR⎵S individuals (resistant to C1 and C19, but susceptible to P20) (Supplementary GWAS Table S3), which can be explained by an epistatic relationship linking the C- and the E-loci. This epistasis confers P20-susceptibility to individuals susceptible to C1 and C19, i.e. presenting the “cc” genotype, regardless of their genotype at the E-locus. This genetic model is presented in Fig. 5 (without variation at the B-locus, see below). In the present study, we mostly considered variation at the C- and E-loci, as they seem to play a major role in the diversity of resistotypes in our study population.

**Figure 5.**
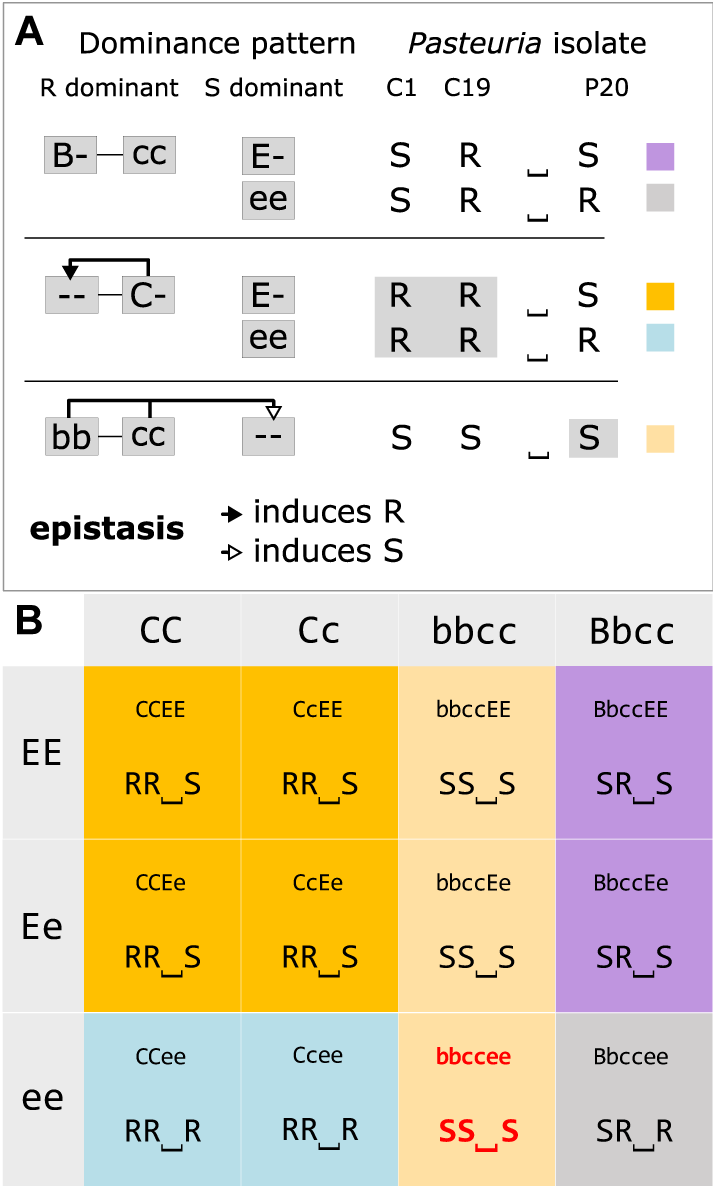
Model for the genetic architecture of resistance to C1, C19 and P20 *Pasteuria ramosa* isolates in the Aegelsee *Daphnia magna* population as inferred from the GWAS analysis (Fig. 4) and the genetic crosses (Tables 1 & 2). **A**: Schematic representation of the genetic model. Resistance to C1 and C19 is determined by the ABC-cluster as described in Metzger et al. (2016), and the model is extended to include the newly discovered E-locus. The dominant allele at the B-locus induces resistance (R) to C19 and susceptibility (S) to C1. The dominant allele at the C-locus confers resistance to both C1 and C19, regardless of the genotype at the B-locus (epistasis). The newly discovered E-locus contributes to determining resistance to P20. Resistance is dominant at the C-locus (resistance to C1 and C19) but recessive at the E-locus (resistance to P20). Homozygosity for the recessive allele at the B- and C-loci induces susceptibility to P20, regardless of the genotype at the E-locus (epistasis). Hence epistasis can only be observed phenotypically in the “bbccee” genotype, which has the resistotype SS⎵S. Without epistasis, the “bbccee” genotype is expected to have the phenotype SS⎵R, a phenotype we never observed in the population or in our genetic crosses. **B**: Multi-locus genotypes and resistotypes at the B-, C- and E-loci. Resistotypes are grouped by background color. As the C-allele epistatically nullifies the effect of the B-locus, only combinations of the B- and E-loci are shown where the C-locus is homozygous for the c-allele. This model does not consider variation at the A-locus, as the recessive allele at this locus is believed to be fixed in the Aegelsee *D. magna* population.

To test the genetic model derived from the GWAS, we investigated segregation of resistotypes among selfed offspring of *D. magna* genotypes with diverse resistotypes. *D. magna* reproduces by cyclical parthenogenesis, in which asexual eggs produce clonal lines and sexual eggs can be used to perform genetic crosses. Our genetic model allowed us to predict the segregation of genotypes and phenotypes, which can then be compared to the observed segregation patterns among selfed offspring. From 24 host genotypes (F0 parent clones), we produced 24 groups of selfed F1 offspring. Twenty-two F0 clones included animals with all possible combinations of alleles at the C- and E-loci, while two F0s showed the rare variation at the B-locus and variation at the E-locus. Expected and observed resistotype frequencies are presented in Tables 1 and 2 and detailed for each F1 group in Supplementary Selfing results Tables S4 to S14. In the 22 F1 groups showing variation at the C- and E-loci, segregation of offspring followed the predictions of our genetic model, i.e. we observed all expected resistotypes and saw no significant deviations from the expected frequencies. These data clearly support the genetic model for resistance at the C- and E-loci.

**Table 1.**
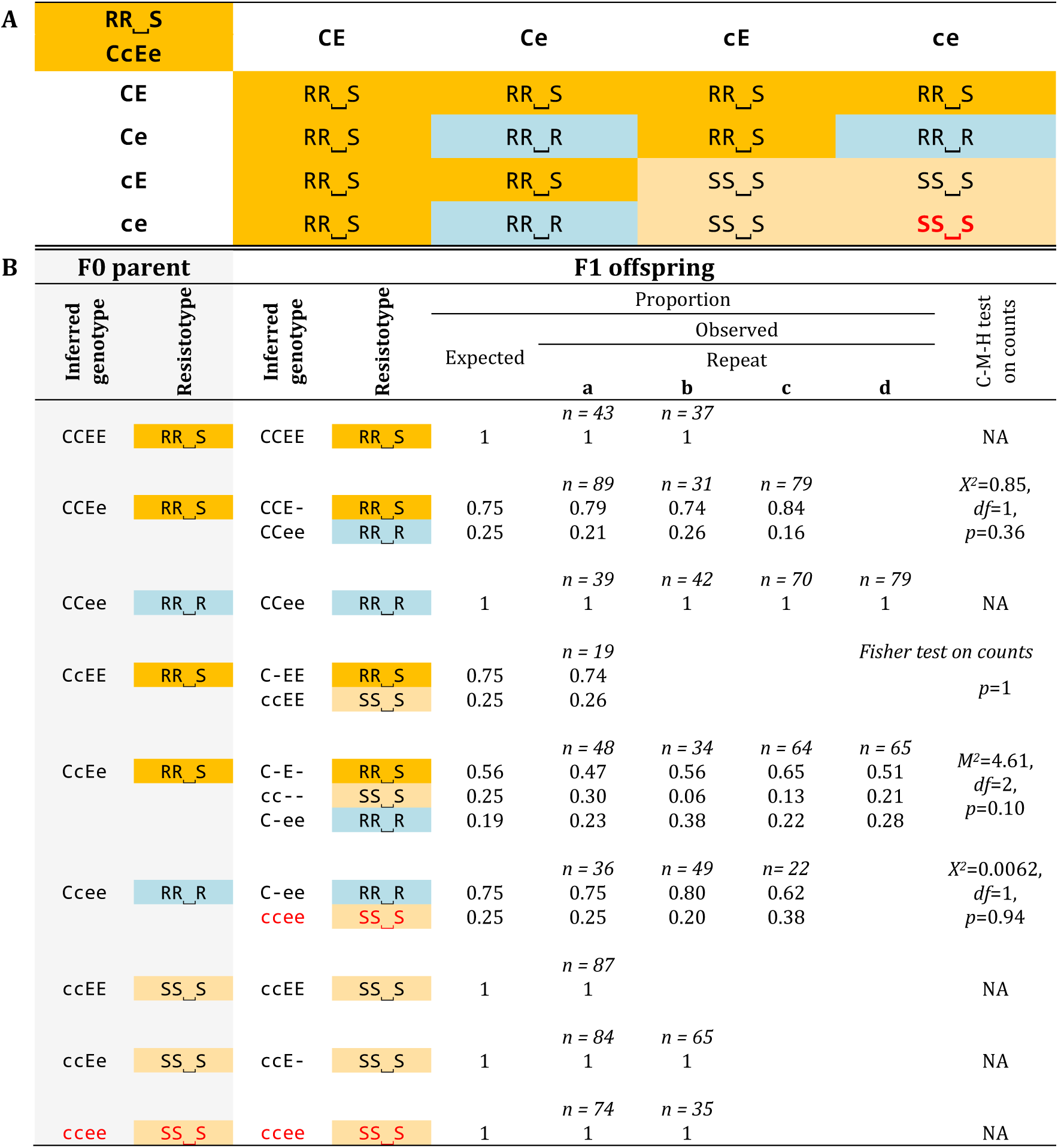
Genetic crosses of resistance phenotypes (resistotypes) from the Aegelsee *Daphnia magna* population. Only the C- and the E-loci are considered. **A**: Punnett square for all possible gamete combinations according to our genetic model of resistance inheritance. The table shows the resistotypes (grouped by background colour) from the 16 combinations of gametes from a double heterozygote for the C- and the E-loci. The bottom right cell (red font, bold) represents offspring individuals where the epistatic interaction between the C- and the E-loci is revealed (Fig. 5). **B**: Results from selfing of *D. magna* clones. Resistotypes of F0 mothers and F1 offspring groups were obtained using the attachment test, and resistance genotypes of F0 parents at the C- and E-loci were inferred from their resistotypes and the segregation patterns of resistotypes in their F1 offspring. Expected resistotype proportions within F1 groups were calculated using the genetic model presented in the Punnett square and the R package “peas” (Fig 5, Supplementary Peas Doc. S1. Detailed results and statistical analyses for each cross are presented in Supplementary Selfing results Tables S4 to S12 and Table S15. Segregation of offspring is presented as proportions, although statistical tests were run on counts. One to four crosses using distinct mother clonal lines (repeats a to d) were conducted for each F0 mother resistance genotype at the C- and E-loci. No variation at the B-locus was observed (all F0 mothers are inferred to have the “bb” genotype according to F1 resistotype segregation).

**Table 2.**
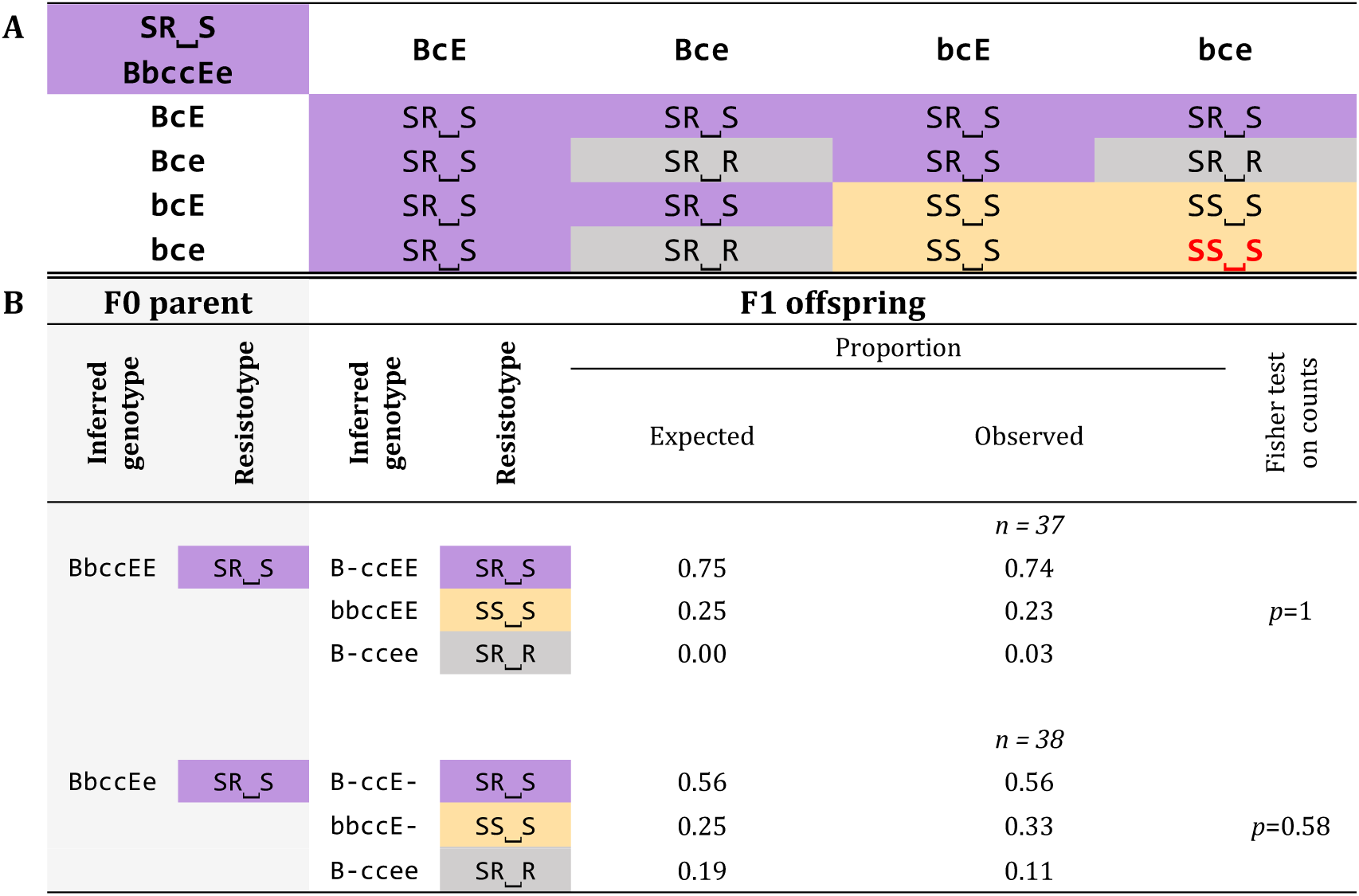
Genetic crosses of resistance phenotypes (resistotypes) from the Aegelsee *Daphnia magna* population considering the B- and the E-loci, with the C-locus fixed for genotype cc. **A**: Punnett square for all possible gamete combinations according to our genetic model of resistance inheritance. The table shows the resistotypes (grouped by background colour) resulting from the 16 combinations of gametes from a double heterozygote for the B- and the E-loci. The bottom right cell (red font, bold) represents offspring individuals where the epistatic interaction between the B-, the C-, and the E-loci is revealed (Fig. 5). **B**: Results from selfing of *D. magna* clones. Resistotypes of F0 parents and F1 offspring were obtained using the attachment test, and resistance genotypes of F0 parents at the B-, C- and E-loci were inferred from their resistotypes and the segregation patterns of resistotypes in their F1 offspring. Expected resistotype proportions of F1 were calculated following the genetic model outlined in the Punnett square and using the R package “peas” (Fig 5, Supplementary Peas Doc. S2). The detailed results and statistical analyses for each cross are presented in Supplementary Selfing results Tables S13 to S15. Segregation of offspring is presented as proportions, although the statistical tests were run on counts.

As described above, earlier research (Metzger et al. 2016; Bento et al. 2017) has shown that the dominant allele at the C-locus interacts epistatically with the A- and B-loci (all are part of the ABC cluster), such that variation at the A- and B-loci becomes neutral when a C-allele is present. We assume that the a-allele is fixed in the Aegelsee *D. magna* population, so that only variation at the B-locus influences the C1- and C19-resistotypes in individuals with the “cc” genotype. As variation at the B-locus is very rare in our *D. magna* study population and could not be included in the GWAS analysis, we selfed two *D. magna* genotypes that presented the very rare SR⎵S resistotype, whose underlying genotype at the ABC-cluster we expect to be “B-cc” (probably “Bbcc”, considering the B-allele is rare in the population). In the F1 offspring of the two F0 parents with the SR⎵S resistotype and the “Bbcc--” genotype, we observed SR⎵R individuals. We speculate that SR⎵R animals have the genotype “B-ccee”, indicating that the epistatic relationship previously described between the C- and the E-loci (“cc” acts epistatically on the E-locus) should also include the B-locus. Thus, “bbcc” acts epistatically on the E-locus (Fig. 5). The two groups of selfed F1 offspring involving “Bb” heterozygotes showed a good fit between this expectation in the expanded model and the observed phenotypic segregation. We observed one SR⎵R offspring produced from a SR⎵S parent with the inferred “aaB-ccEE” genotype, which is not expected in our model (Table 2B, lower panel), but typing mistakes cannot be fully ruled out.

#### Linking the genomic regions and the genetic model of resistance

To verify whether the segregation of the genomic regions we discovered in our GWAS determined the phenotypic segregation pattern in our crosses, we designed pairs of size-polymorphic markers in the genomic regions of the C- and the E-loci and tested whether variation at the genetic markers was associated with resistotype variation. Of our four markers, DMPR1 (C-locus) and DMPR3 (E-locus) showed better linkage with their respective resistance loci (99.6% and 94.8% match, respectively) compared to DMPR2 (C-locus) and DMPR4 (E-locus) (91.4% and 69.4% match, respectively) (Supplementary Markers linkage Tables S16 to S18). We thus further based our scoring of resistance genotypes on the marker genotypes of DMPR1 and DMPR3. In 20 F1 groups representing all possible combinations of alleles at the C- and E-loci, the segregation of marker genotypes in the F1 offspring followed our genetic model predictions, i.e. all expected genotypes were observed, with no statistically significant deviations from the expected frequencies. In two F1 groups from “CCEe” and “CcEe” F0 parents, the E-locus markers appeared not to be linked to the E-locus. Genotype markers had assigned the “EE” genotype to the F0 parent and F1 offspring, but phenotypic segregation in the F1 offspring indicated the parent should have the “Ee” genotype. These two groups were not included in the statistical analysis (Supplementary Selfing results Tables S5 and S8). Together, these results show that the genomic regions found in the GWAS are indeed associated with the segregation of resistotype in the F1 selfed offspring, supporting the genetic model of resistance derived from the GWAS and experimental selfing.

### Allele frequency fluctuations caused by parasite-mediated selection

To represent the genetic change associated with the evolution of resistance in our study population, we inferred the dynamics of resistance genotypes and allele frequencies over the two-year study period using (i) observed longitudinal resistotype data (Fig. 1), (ii) the genetic model of resistance yielded by the present study (Fig. 5) and (iii) allele polymorphism at the C- and E-loci in a spring sample of the host population (Supplementary Allele frequency Fig. S4 and Table S20). In this spring sample (n = 108), the C- and E-loci were in Hardy-Weinberg equilibrium (*f(C-locus)* = 0.514, Fisher test: *p* = 1 and *f(E-locus)* = 0.259, Fisher test: *p* = 1) (Supplementary Allele frequency Table S20). We used resistance genotype frequencies within resistotypes in this sample to infer the evolution of genotype and allele frequencies over time in the population and applied these genotype frequencies to the longitudinal resistotype frequency data (Fig. 1), assuming only clonal reproduction in the population during the active season and Hardy-Weinberg equilibrium. We acknowledge that, being under selection, the population would no longer be in Hardy-Weinberg equilibrium during the epidemics. As expected, we observed an increase in the frequency of resistant alleles (C- and e-alleles) over the course of the epidemics due to parasite-mediated selection. The change in allele frequencies, however, appeared small compared to the strong change in resistotype and genotype frequencies (compare Fig. 6A and B).

**Figure 6.**
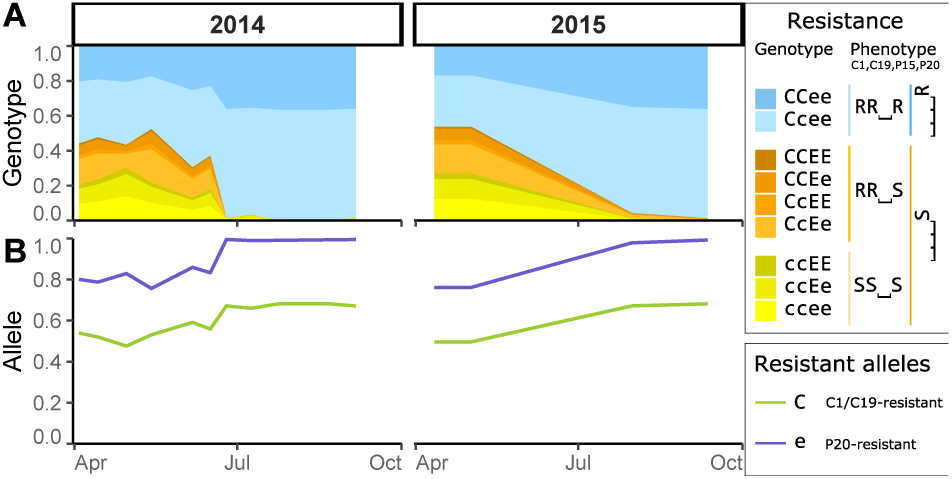
Dynamics of parasite-mediated selection in the Aegelsee *Daphnia magna* population. Using resistotype (resistance phenotype) frequency over time (Fig. 1) and genotype frequency from a sample in spring 2015 (Supplementary Allele frequency Table S20), we infer genotype and allele frequency over time. We assume only clonal reproduction during the active season of *D. magna* and Hardy-Weinberg Equilibrium (HWE) in the population over time, although we acknowledge that, in reality, selection will make the population deviate from HWE. **A**: Genotype/resistotype frequencies plotted across time during the study period. Genotypes that did not match the expected resistotype were excluded (see Supplementary Allele frequency Table S19, Fig. S4 and Table S20). **B**: Allele frequency of the C- and the E-loci over time. The alleles that confer resistance to the host (C- and e-alleles) increased in frequency during the epidemics.

## Discussion

This present study aims to assess how annual epidemics by a parasitic bacterium, *Pasteuria ramosa*, influence resistance and its underlying genes in a natural host population of the crustacean *Daphnia magna*. Over the course of epidemics in two consecutive years, we observed drastic changes in resistance phenotype (resistotype). Using experimental infections and fitness measurements on wild-caught individuals, we showed that these changes in resistotype frequency were caused by a local parasite common during the early phases of the epidemics. A genome-wide association study (GWAS) and laboratory crosses enabled us to find the resistance genes that responded to this selection and their mode of inheritance. We pinpointed the genetic architecture of resistance to two genomic regions with dominance and epistasis, thus bridging the gap between natural selection on phenotypes and the underlying genetic response.

### Parasite-mediated selection

Over the two consecutive years of this study, resistotype frequencies in the host population changed drastically during the parasite epidemics, but remained stable outside of the epidemics (Fig. 1)—a pattern consistent with the host population being under strong selection for resistance to *P. ramosa*. The P20 *P. ramosa* isolate, collected during the early epidemic, revealed that this local parasite plays a major role in the resistotype frequency changes: P20-susceptible individuals are much more susceptible to the local parasite than P20-resistant individuals in experimental infections (Fig. 2A). Infected P20-susceptible genotypes also become infected earlier and produce fewer offspring than P20-resistant individuals (Fig. 2B & C), revealing a stronger fitness impact of infection by the local parasite. Field data confirmed this result, as wild-caught P20-susceptible individuals are infected more frequently and produce fewer offspring than infected P20-resistant individuals (Fig. 3), again showing the higher virulence of the parasite in these P20-susceptible individuals. Together, these results reveal strong and rapid parasite-mediated selection on host resistotypes in the Aegelsee *D. magna* population.

Although the parasite P20 was isolated during the early phase of the yearly epidemics, previous research also shows other parasite genotypes in the Aegelsee population (Andras and Ebert 2013) that, as we observed in an earlier year, become more common in infected hosts later in the epidemics (Supplementary *Pasteuria* Fig. S1). We speculate that these later-season isolates may represent different parasite infectotypes (infection phenotypes). Consistent with this, we observed that animals resistant to P20 did, in fact, become infected, both in the field and in the laboratory (Fig. 1, 2 and 3). This present study focuses on natural selection during the early part of the epidemics, which, as our data and data from other years shows, has a fairly consistent selection pattern (Ameline et al. in prep), being mainly defined by a drastic increase in P20-resistant individuals from around 50% to almost 100% within a period of two to three months (Fig. 1).

The yearly *P. ramosa* epidemics in our population are typical during the summer, when water temperature rises above 15 °C (Ameline et al. in prep). It has been suggested that parasite-mediated selection in the *D. magna - P. ramosa* system is strongest at 20-25 °C (Mitchell et al. 2005; Vale et al. 2008). Given climate change model predictions of pond warming and longer seasons with temperatures above 15 °C, selection for resistance can thus be expected to intensify in our study population. Warming could also affect the evolution of stress tolerance, as exposure to the pathogen disrupts the host’s ability to cope with thermal stress in this system (Hector et al. 2019). Environmental factors may also change the mode of selection: it has been shown that, under some temperature and food availability conditions, hosts in this system become more tolerant, thus potentially increasing parasite prevalence and slowing down coevolution (Vale et al. 2011). Parasite fitness may also be influenced by the interaction of genotype and environmental factors such as temperature and food availability (Vale and Little 2009, in a plant-parasite system: Laine 2007). Thus, while natural selection on resistance is precipitated on a high specificity of host-parasite interactions in the *D. magna* - *P. ramosa* system, it may also be linked to environmental conditions.

The composition of the resistotypes at the beginning of the two seasons (2014 and 2015) in which we monitored this system was strikingly similar, which is surprising given that selection increased resistance over the course of the summer 2014. While this question is not part of the current study, there are a few tentative explanations for this observation. First, resting eggs, which form the basis of the new population in the following year, are produced as early as mid-June before selection has eradicated some of the resistotypes. Second, epistasis and dominance can protect alleles from natural selection, thus slowing down the response to selection (Feldman et al. 1975). Our study, as well as earlier studies on this system (Luijckx et al. 2012; Luijckx et al. 2013; Metzger et al. 2016), all indicate strong epistasis and dominance. Further studies are needed to understand how much resting egg production and the genetic architecture of resistance explain the slow response of selection observed across seasons in the Aegelsee *D. magna* population.

### Genetic architecture of resistance

To understand the genetic architecture of resistance loci under selection in our study population, we combined a GWAS using *D. magna* clones with segregating resistotypes together with a series of genetic crosses. We found that host resistance to the bacteria is determined by a previously described supergene, the PR-locus containing the ABC-cluster (Bento et al. 2017), and by a newly discovered locus on a different chromosome, the E-locus (Fig. 4). Taken alone and in the right genetic background, i.e. when there is no epistatic relationship, each of these two loci show Mendelian segregation with resistance being dominant (C-locus) or recessive (E-locus) (Supplementary GWAS Fig. S3). The two loci interact epistatically with each other, resulting in a complex pattern of inheritance (Fig. 5).

As resistance genes are hypothesized to maintain diversity, notably through balancing selection (Llaurens et al. 2017; Wittmann et al. 2017; Connallon and Chenoweth 2019), they are often found to have different dominance patterns and epistatic interactions (Saavedra-Rodriguez et al. 2008; González et al. 2015; Conlon et al. 2018), In our study population, we found resistance polymorphisms mainly at the C- and E-loci, which produced the three common resistotypes RR⎵R, RR⎵S and SS⎵S. The first two letters (C1- and C19-resistotypes) were mainly influenced by the C-locus, and the last letter (P20-resistotype) by a combination of the C- and E-loci (Fig. 5).

In the *D. magna - P. ramosa* system, the ABC-cluster has been shown to play a major role in host resistance and the evolutionary dynamics of resistance (Routtu and Ebert 2015; Bento et al. 2017; Bourgeois et al. 2017). Our results confirm the role of this cluster in a natural population and describe a new resistance region in the *D. magna* genome that is polymorphic in the Aegelsee population. Multi-locus polymorphisms have been shown to underlie parasite resistance in host-parasite coevolution (Sasaki 2000; Tellier and Brown 2007; Cerqueira et al. 2017). In the Aegelsee *D. magna* population, there seems to be no variation at the A-locus and little at the B-locus. The observed variation at the B- and C-loci is consistent with the genetic model of resistance at the ABC-cluster described in Metzger et al. (2016). Also, resistance to *P. ramosa* isolate P15 (influenced by the D-locus, Bento et al. submitted) remains fairly consistent, with the vast majority of animals being susceptible to P15 (Fig. 1). Resistance to *P. ramosa* P21, isolated from our study population, varies only towards the end of the summer epidemic (Ameline et al. in prep). In summary, resistance to P20 plays a major role in the early epidemics and most resistotype diversity we measured in the Aegelsee *D. magna* population is well explained by genotypic variation at the C- and the E-loci. We do not expect other resistance regions to hold similar influence in the early phase of the epidemic, although other *P. ramosa* isolates and their associated resistance regions in the *D. magna* genome may gain importance later in the epidemic in the Aegelsee population. The study of these parasite genotypes may reveal further resistance loci and help explain late season resistance variation in this population.

Resistance segregation in *D. magna* is currently best explained by a genetic model where each locus contains just two alleles. This model was compiled by studies that used either mapping panels created from a few *D. magna* genotypes or, as here, host genotypes from one focal population. Additional resistance alleles may be revealed instead of new resistance regions if we test the genetic model on a larger panel of host and parasite genotypes.

The E-locus, which codes for resistance to P20, is situated on linkage group (LG) 5 (genome version 2.4: Routtu et al. 2014) and appears as a large region of 3.1 Mb (Fig. 4). In this regard, the E-locus is similar to the ABC-cluster, a well-characterized, non-recombining and extremely divergent region on LG 3 (Bento et al. 2017). We hypothesize that non-recombining genomic structures, i.e. supergenes, could facilitate adaptation via association of advantageous alleles in host–parasite coevolution (Joron et al. 2011; Llaurens et al. 2017). Such large, diverse genomic regions are difficult to study because the absence of recombination hampers fine mapping (Bento et al. 2017). Therefore, we do not know where the actual resistance loci lie within the ABC- and E-loci regions. Within the E-locus region, we find four sugar transferases. Glycosylation genes are candidates to explain variation of resistance in this system (Bento et al. 2017; Bourgeois et al. 2017).

We created 22 F1 offspring groups from the three common resistotypes in our study population. Segregation of resistance phenotypes and genotypes among the selfed F1 strongly supported the genetic model of resistance, consisting of the C- and E-loci and their epistatic interaction, produced by the GWAS (Tables 1 & 2, Supplementary Selfing results Tables S4 to S15). Two F1 offspring groups showed rare variation at the B-locus, suggesting yet an additional epistatic interaction in this model besides the previously described role of the B-locus for the *P. ramosa* C1- and C19-resistotypes. This consisted of the “bbcc” genotype that causes susceptibility to P20, irrespective of the genotype at the E-locus (Fig. 5). However, this modified model needs to be further investigated and verified with more genetic crosses. The overall good fit between the expected and observed segregation of genetic markers in the F1 offspring groups shows that the associated genomic regions discovered by the GWAS are indeed responsible for the modes of inheritance seen in the genetic crosses. Thus, our study uncovers the mode of inheritance and underlying genomic regions of resistance phenotypes that evolve under parasite-mediated selection in the Aegelsee *D. magna* population.

## Conclusion

In this study, we demonstrate rapid parasite-mediated selection in a natural plankton population. We find the genomic regions associated with resistance under selection and describe their mode of inheritance. This knowledge will allow us to conduct direct measurements of resistance allele frequency over larger timescales and to test theories on the dynamics of host and parasite evolution, for example by tracing genetic changes in the resting stages of *Daphnia magna* derived from the layered sediments in ponds and lakes (Decaestecker et al. 2007). Pinpointing resistance loci can also be used to infer mechanisms of selection in the host with the molecular evolution tool box (Charlesworth 2006; Fijarczyk and Babik 2015; Hahn 2018). Our model of resistance consists of a few loci linked with epistasis and different dominance patterns, characteristics that have been shown to be relevant in coevolution, particular when balancing selection maintains diversity at resistance genes (Tellier and Brown 2007; Engelstädter 2015; Conlon et al. 2018). The genomic regions we pinpoint can now be further studied using the molecular evolution tool box, e.g. testing for genomic signatures for balancing selection (Charlesworth 2006). Hence, a precise knowledge of the genetic architecture of resistance opens the door to addressing wider evolutionary questions. For example, the Red Queen theory states that host-parasite interactions may explain the ubiquity of sex and recombination (Salathé et al. 2008). We further emphasize the importance of context-dependent parasitism (Vale et al. 2011) in shaping the tempo and mode of evolution. Understanding the impact of parasite-mediated selection on a host population at the genetic level will lead to further studies on the effect of environmental changes on epidemics (Auld and Brand 2017).

## Material and methods

### Study site

Our study site is the Aegelsee, a pond near Frauenfeld, Switzerland (code: CH-H for Hohliberg; coordinates: 47.557769 N, 8.862783 E, about 30000 m^2^ surface area) where *D. magna* is estimated to have a census population size over ten million individuals and an overwintering resting egg bank of about the same size. In early fall, the pond is used as a waste repository by a sugar factory: they progressively lower the water level from May to September and from October, warm ammoniacal condensation water is released into the pond, warming the water temporarily to 40-60 °C and killing all zooplankton, but not the resting eggs. In winter the pond usually freezes over, and in April, *Daphnia* and other invertebrates hatch from resting eggs. We sampled the pond in February 2014 and March 2015 and did not find any *Daphnia*, suggesting little or no overwintering of planktonic animals. Besides *D. magna*, the plankton community includes *D. pulex, D. curvirostis* and a diverse array of other invertebrates, among them copepods, ostracods, rotifers and corixids. The *D. magna* population experiences strong yearly epidemics of *P. ramosa*, reaching prevalence of 70-95%. Infections by other parasites were only rarely observed. The other *Daphnia* species in the pond were never observed to be infected by *P. ramosa*.

### Temporal monitoring

In 2014 and 2015, we sampled the host population every two to three weeks from early April to early October to study the impact of the pathogen epidemics. For each sampling date, we aimed to obtain about 100 cloned host lines (produced as iso-female lines). To achieve this, we randomly took about 200-300 female *D. magna* from the sample, placed them in 80-mL jars filled with ADaM (Artificial *Daphnia* Medium, Klüttgen et al. 1994, as modified by Ebert 1998) and let them reproduce asexually. Oversampling was necessary during the hot summer months, as many animals would die for unknown reasons within 48 hours under laboratory conditions. This mortality was, to the best of our knowledge, not disease related. Over the following three weeks, we screened animals for *P. ramosa* infections by checking for the typical signs of disease: gigantism, reddish-brownish opaque body coloration and empty brood pouch. Infected animals that had not yet reproduced asexually were treated with tetracycline (50 mg.L^-1^) (an antibiotic which kills Gram-positive bacteria) until an asexual clutch was observed, usually after about two weeks. They were fed 25 million cells of the unicellular green algae *Scenedesmus* sp. three times a week, and the medium was renewed every two weeks, although feeding and fresh medium protocols were adapted according to the size and number of animals in a jar.

### Resistotype assessment: the attachment test

We estimated resistance phenotype (resistotype) frequencies in the samples of cloned hosts using four *P. ramosa* isolates (C1, C19, P15 and P20). We isolated the parasite, P20, from our study population at the start of the epidemic on 13 May 2011 and subsequently passaged it three times through a susceptible *D. magna* host clone from the same population. The three other *P. ramosa* clones or isolates had been previously established in the laboratory: C1 (clone), originated from a *D. magna* population in Russia (Moscow), C19 (clone) from Germany (Gaarzerfeld) and P15 (isolate) from Belgium (Heverlee) (Bento et al. submitted; Luijckx et al. 2011). We used these three *P. ramosa* allopatric isolates in the present study to implement our working genetic model for resistance (Luijckx et al. 2012; Luijckx et al. 2013; Metzger et al. 2016). Spore production in the laboratory followed the protocol by Luijckx et al. (2011).

The resistotypes of *D. magna* clones were assessed using a spore attachment test (Duneau et al. 2011). Bacterial spores attach to the foregut or the hindgut of susceptible host clones, with the absence of attachment indicating host resistance. We exposed each individual *Daphnia* to 8000 (C1, C19) or 10000 (P15, P20) fluorescent spores following the protocol of Duneau et al. (2011). We used higher spore doses for P15 and P20 because previous observations had shown that fewer of these isolate spores attach to the host’s oesophagus, resulting in a weaker fluorescent signal. For each host genotype–parasite isolate combination, we used three replicates—or more when attachment phenotype was not clear. A clone was considered susceptible to the bacterial isolate when more than half of its replicates showed clear attachment. Its overall resistotype is the combination of its resistotypes to the four individual *P. ramosa* isolates in the following order: C1, C19, P15 and P20, e.g. a clone susceptible to all four isolates would have resistotype SSSS. Since resistance to P15 had low variability in our study population, this isolate was only considered in one of the experiments presented here and was otherwise represented with the placeholder “⎵”, e.g. “RR⎵R resistotype”. Overall, we observed that the attachment test for P20 necessitated more repeats than for the other isolates to yield a clear and consistent result (Three repeats were used for C1, C19 and P15 whereas six to nine repeats were often needed for P20).

### Experimental infections of resistotypes

As an initial assessment of the parasite’s fitness impact on the host population, we conducted experimental infections on a representative sample of the spring 2014 host population.

#### Sampling of host and parasite populations

We collected surface sediment from five different points in the pond in February 2014, before onset of the natural hatching season and placed one hundred *D. magna* ephippia from each replicate in 80-L containers with 30 L ADaM. The five containers were placed outdoors under direct sunlight and checked for hatchlings every two days. We recorded hatching dates and cloned hatchlings in the laboratory where we then scored their resistotypes.

For the infection experiment, we used parasites collected from the ongoing epidemic in the pond. We collected 20 randomly chosen infected individuals during the first phase of the epidemic in early June 2014. These field-infected animals were kept in the laboratory under ad libitum feeding conditions. Shortly before their expected death, we pooled all 20 animals, homogenized them to produce a spore suspension and froze it at −20 °C. A placebo suspension was produced from 20 homogenized uninfected *D. magna*. This spore mixture was not passaged before we used it, so, in contrast to the isolates used for the attachment test, it represents a population sample of the parasite.

#### Experimental infections

Among the four predominant resistotypes we observed in the cloned cohort of spring hatchlings (SSSS, RRSS, RRSR and RRRR), we used 20 clones each from the more common resistotypes SSSS, RRSS and RRSR and ten of the less common resistotype RRRR for an infection experiment. From each of these 70 clones, we produced five replicate lines, and these 350 lines were maintained individually in 80-mL jars. To reduce maternal effects before the experiment, we kept all lines for three generations in the same experimental conditions: 20 °C, 16:8 light:dark cycle, ADaM medium and daily ad libitum feeding of 8 million *Scenedesmus* sp. cells per jar. The three generations were produced as follows: as soon as a female produced a clutch, she was discarded. When the offspring were mature, a single female was kept in the jar until she in turn produced a clutch. The medium was changed every four days or when the females released offspring. We exposed two-to three-day old juveniles from all replicates to the parasite spore suspension by placing individual *D. magna* in 10 mL of medium with 10000 spores. Additionally, three controls from the third-generation offspring were randomly taken from among the five replicates for each clone (n = 210) and were exposed to the equivalent volume of placebo suspension. Three days after exposure, the jars were filled to 80 mL. Medium was changed after ten days, and then every four days until the end of the experiment. Jars were monitored daily for 35 days. We recorded infection occurrence, clutch number and time of infection (when visible signs of infection were observed). Controls did not get infected and produced offspring at regular intervals.

#### Statistical analysis

We tested the effect of the full resistotype or P20-resistotype on the three dependent variables: infection (binary: 1/0), clutch number (integer) and time of infection (continuous). Replicates were nested within clones, which were nested within full resistotypes or P20 resistotype. We fitted general linear models using binomial data family type for infection and quasi-Poisson for clutch number and time to infection. For clutch size and time to infection, only data on infected individuals was used.

### Infection phenotypes of field-collected hosts

As a second assessment of the impact of the local parasite on the host population, we measured fitness traits of animals caught during the epidemics. Because the infection experiment described above (carried out in the previous year) indicated that P20 played a strong role, we focused on this parasite isolate. On June 7 and 28, 2015, we collected large *D. magna* samples from our study site and measured body length, from the top of the head through the eye to the base of the tail spine. We kept all females (n = 331) individually under ad libitum feeding conditions, each in about 80 mL medium. We recorded clutches (time and size) and the onset time of disease symptoms over the following three weeks. After parasitic castration was evident, we cured animals with tetracycline. These data have also been reported in a paper describing the disease phenotype under natural conditions (Savola and Ebert 2019).

Using generalized linear models, we tested the effect of the P20-resistotype on infection and fecundity, taking body size into account. Sampling date was included as a fixed effect since there are only two sampling dates. Interaction terms were excluded from the model when not significant (p > 0.1). We fitted a general linear model using quasibinomial data family type for infection, and a negative binomial generalized linear model for total fecundity (R packages used: MASS: Venables and Ripley 2002, lme4: Bates et al. 2015).

### Genome-wide association study

Because our experiments revealed that resistance to P20 plays a major role in the disease dynamics in both laboratory experiments and the field, we used a genome-wide association approach to investigate the genetic architecture of resistance to this local parasite with thirty-seven clones that presented the three most common resistotypes in our study population, excluding P15-resistotype (16 RR⎵R, 10 RR⎵S and 11 SS⎵S). All clones were derived directly from our study population (Supplementary GWAS Table S3).

#### Whole-genome DNA extraction, sequencing and bioinformatics

To remove microbial DNA, individuals were treated for 72 h with three antibiotics (streptomycin, tetracycline, ampicillin at a concentration of 50 mg.L^-1^ each in filtered water) and fed twice daily with 200 µL of a dextran bead solution (Sephadex G-25 Superfine by Sigma Aldrich: 20-50 µm diameter at a concentration of 5 g.L^-1^) to remove algae from the gut. DNA was extracted from 15-20 adult animals using an isopropanol precipitation protocol (QIAGEN DNeasy Blood & Tissue Kit). Paired-end 125-cycle sequencing was performed on an Illumina HiSeq 2000.

Raw reads were aligned using BWA MEM (Li and Durbin 2009) on the *Daphnia magna* draft genome (v.2.4) and a genetic map (Routtu et al. 2014). BAM alignment files were then filtered for quality, and PCR duplicates were removed using PICARD tools (http://broadinstitute.github.io/picard/). Variant calling was performed using freebayes (v. 0.9.15-1). VCF files were then filtered using VCFTOOLS v. 0.1.12b (Danecek et al. 2011) to include SNPs with a minimum quality of 20, a minimum genotype quality of 30, and a mean sequencing depth between 10X and 50X. Only SNPs that passed filters in every sample were included in subsequent analyses, resulting in a dataset of 510,087 SNPs. Association analyses were performed using the command “-assoc” in PLINK (Purcell et al. 2007), which compares allele counts between cases and controls and outputs a p-value from a chi-square test with one degree of freedom. Five pairwise comparisons were performed to identify possible candidates for resistance to C1, C19 and P20: (i) SS⎵⎵ vs. RR⎵⎵, (ii) SS⎵S vs. RR⎵S, (iii) ⎵⎵⎵S vs. ⎵⎵⎵R, (iv) RR⎵S vs. RR⎵R and (v) SS⎵S vs. RR⎵R. We corrected for the genomic inflation of p-values (λ) that may have resulted from relatedness between samples using the R package GenABEL (Aulchenko et al. 2007). Lambda was calculated excluding SNPs from linkage groups 3 and 5, since these scaffolds displayed an excess of strongly associated markers. We divided raw chi-square scores by λ to obtain corrected p-values using R commands “pchisq” and “estlambda”.

For each SNP:

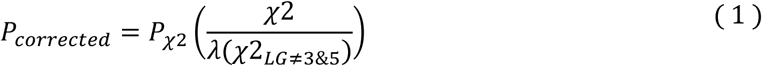

Histograms of corrected p-values were examined to confirm their uniform distribution. We estimated the minimum false discovery rate incurred when a given p-value was identified as significant (so-called q-value) from the set of corrected p-values using the R package “qvalue” (Storey et al. 2015).

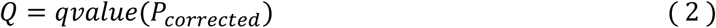

The minimum significant threshold for a given association was then calculated as the maximum corrected p-value with a q-value less than 5%.

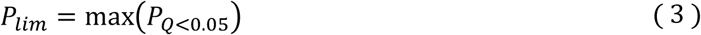

The “gg.manhattan” function in R was used to display manhattan plots of the comparisons between different resistotypes (https://github.com/timknut/gg.manhattan/). We used BEDTOOLS (v 2.25.0) to extract genes found in the associated candidate regions, using the 2011 annotation of the genome (available at: wfleabase.org).

### Assessment of resistotype segregation

The genetic model that resulted from the GWAS analysis allowed us to make predictions about the segregation of resistotypes in sexually reproducing *D. magna* lines. To test these predictions, we selfed *D. magna* clones with different resistotypes. Selfing is possible with *D. magna* because the same clonal line can produce sons (asexual production) as well as eggs by sexual production. The latter need fertilization by males. The resulting sexual eggs must undergo an obligatory resting phase before they can hatch (Ślusarczyk et al. 2019). The resistotypes of the selfed offspring (F1) were examined to assess whether their segregation matched expectations from the genetic model derived from the GWAS.

All clones used for the genetic crosses derived from the study population. We selfed five to ten *D. magna* clones of the three common resistotypes (RR⎵R, RR⎵S and SS⎵S) and two clones of a rare resistotype (SR⎵S), following the protocol from (Luijckx et al. 2012). Hatching of selfed offspring is not always successful, resulting in uneven sample sizes. We obtained between 19 to 89 selfed offspring from each of 22 parent clones (Supplementary Selfing methods Table S21). Their resistotypes were assessed with the attachment test. Samples from each clonal line were stored at −20 °C for future DNA extraction and genotyping.

#### Predictions of segregation patterns

We compared the resistotype segregation patterns in the selfed offspring to predictions in our genetic model. To calculate proportions of expected phenotypes, we developed an R package called “peas” (https://github.com/JanEngelstaedter/peas) that enables the user to predict distributions of offspring genotypes and phenotypes in complex genetic models with Mendelian inheritance (Supplementary Peas Docs. S1 and S2). We compared these predictions to the segregation patterns from our selfed offspring using the Cochran-Mantel-Haenszel (C-M-H) test for repeated tests of independence. The C-M-H test is applied either to 2×2 tables and outputs a Chi-square statistic (χ^2^) or to larger tables (generalized C-M-H test), where it outputs a M^2^ statistic. When there was only one repeat per parent genotype, we used the Fisher test. When there was only one category of expected and observed phenotype (i.e. no segregation), no test was possible, and expectation and observation showed a perfect match. Following each C-M-H test, assumption of homogeneity of the odds ratio across repeats was confirmed using a Breslow-Day test (R package DescTools: Signorell et al. 2018). However, this test can only be used with 2×2 tables. We ran a Fisher test of independence on each comparison (expected vs. observed for each repeat, Bonferroni corrected) to detect differences in opposite directions across repeats, which would have resulted in a non-significant C-M-H test, but no such differences in direction were detected (see Supplementary Selfing results Table S15 for detailed results of statistical analyses). Tests were run on counts, but for better illustration we present here segregation of offspring as proportions.

### Linking the phenotype to the genotype

We designed PCR-based diagnostic markers physically linked to the resistance loci that the GWAS identified (DMPR1 to 4 for “*Daphnia magna* - *Pasteuria ramosa*” markers, Supplementary GWAS Markers Table S23) and tested if these markers (and their corresponding resistance regions) are indeed associated with the resistotypes, by comparing expected and observed association between marker genotypes and resistotypes (Supplementary Markers linkage Tables S16 to S18). We then used these markers to confirm genotyping of the selfed parents.

#### DNA extraction and PCR-based markers analysis

DNA of parents and selfed offspring was extracted on 96-well PCR plates using a 10-% Chelex bead solution (Bio-Rad) adapted from Walsh et al. (1991). First, individuals were crushed in the wells with 20 µL of deionized water using a customized rack of metallic pestles. We added 150 µL of 10-% Chelex solution and 10 µL of proteinase K and incubated samples for two hours at 55 °C followed by 10 min at 99 °C. Fragment amplification, genotyping and allele scoring was done following the protocol described in Cabalzar et al. (2019) (see Supplementary Markers Table S22 for PCR reaction details).

#### Genotype and allele frequency over time

In a sample from spring 2015 (n = 108), we assessed C- and E-loci genotype proportion in the *D. magna* resistotype groups. *D. magna* individuals were genotyped using the genetic markers DMPR1 and 3, and phenotyped using the attachment test. We applied genotype proportions within resistotype to the longitudinal resistotype frequency data to infer genotype and allele frequency over time in the *D. magna* population.

### Statistical software

Unless otherwise stated, all statistical analyses and graphics were performed using R software version 3.6.1 (R Core Team 2019). Mean values are presented with standard error: mean ± se (Package RVAideMemoire v. 09-45-2, Hervé 2015). Packages used in R for package installation, data manipulation and graphics are the following: package development, documentation and installation: devtools v. 2.2.1 (Wickham, Hester, et al. 2019) and roxygen2 v. 6.1.1 (Wickham et al. 2018), data manipulation: dplyr v. 0.8.3 (Wickham, François, et al. 2019), tidyr v. 1.0.0 (Wickham and Henry 2019), tidyquant v. 0.5.8 (Dancho and Vaughan 2019), tidyverse v. 1.2.1 (Wickham 2017), xlsx v. 0.6.1 (Dragulescu and Arendt 2018), graphics: ggplot2 v. 3.3.0 (Wickham 2016), extrafont v. 0.17 (Chang 2014), scales v. 1.0.0 (Wickham 2018), cowplot v. 1.0.0 (Wilke 2019), gridExtra v. 2.3 (Auguie 2017), ggpubr v. 0.2.3 (Kassambara 2019), ggplotify v. 0.0.4 (Yu 2019), magick v. 2.2 (Ooms 2019), egg v. 0.4.5 (Auguie 2019), ggsci v. 2.9 (Xiao 2018) and png v. 0.1-7 (Urbanek 2013).

## Acknowledgments

We thank Jürgen Hottinger, Urs Stiefel, Kristina Müller, Michelle Krebs, Samuel Pichon, Dita Visozo and Jelena Rakov for help in the field and laboratory. Sequencing for the GWAS analysis was performed at the Genomics Facility at the Department of Biosystem Science and Engineering (D-BSSE, ETH) in Basel. The AB3130xl sequencer used for the markers analysis was operated by Nicolas Boileau. Members of the Ebert group provided valuable feedback on the study and the manuscript. Suzanne Zweizig improved the language of the manuscript. This work was funded by the Swiss National Science Foundation (SNSF) grant No 310030B_166677, the Freiwillige Akademische Gesellschaft (FAG) Basel and the University of Basel. JE was supported by the Australian Research Council through a Future Fellowship (FT140100907).

## Authors contribution

DE and JA designed the overall study. FV and DE designed the infection experiment. FV conducted the infection experiment and analyzed the data. ES and DE designed the fitness measurements. ES conducted the fitness measurements and analyzed the data. YB and DE designed the GWAS analysis. YB conducted the GWAS analysis. CA and DE designed the crossings. CA conducted the crossings and analyzed the data. JE developed the “peas” R package. CA analyzed the data, wrote the manuscript and designed the figures. All authors reviewed the manuscript.

