## Supplementary material for "A two-locus system with strong epistasis underlies rapid parasite-mediated evolution of host resistance"

#### Supplementary *Pasteuria*

*Pasteuria ramosa* genotype frequency over time during the 2011 active season of *Daphnia magna*. Infected animals were collected throughout the season, and the genotype of the parasite was determined using microsatellite markers following the protocol described in (Andras and Ebert 2013).

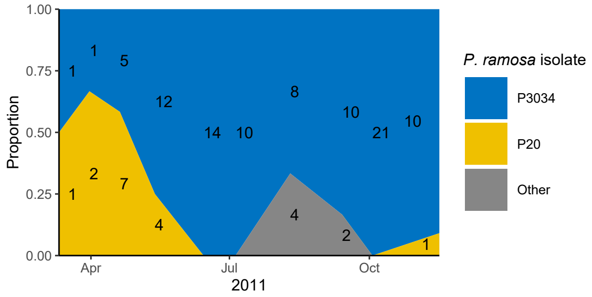

**Figure S1** *Pasteuria ramosa* genotypes frequency over time during the 2011 active season of *Daphnia magna.* A total of 113 infected *D. magna* individuals were sampled throughout the season and their parasite genotype was assessed. The P20 genotype represents about 50% of the parasite diversity among infected hosts at the beginning of the season, a proportion that then drops off during the epidemics. The P3034 genotype represents the most diversity of the parasite population among infected hosts but was not used in this study to score host resistotype (resistance phenotype). Numbers represent the amount of *D. magna* individuals sampled on each collection date.

#### Supplementary Model

The ABC genetic model of resistance in the *Daphnia magna* - *Pasteuria ramosa* host-parasite system. Figure adapted from (Metzger et al. 2016).

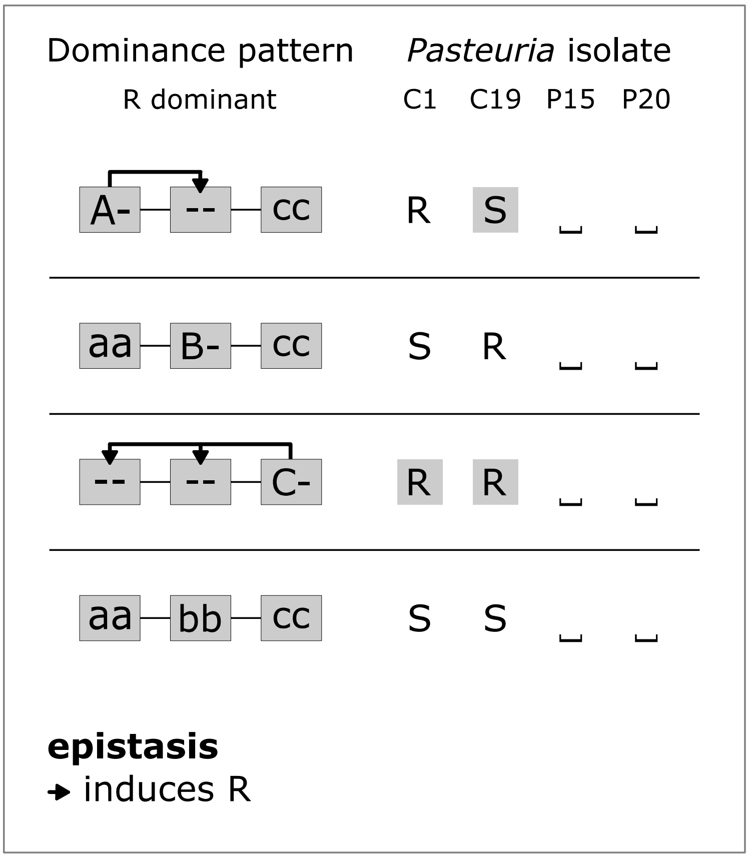

**Figure S2** Genetic model of resistance inheritance at the ABC-cluster in the *D. magna* - *P. ramosa* system. Resistance is dominant at the A-, B- and C-loci. The dominant allele at the A-locus confers resistance to C1 and susceptibility to C19, regardless of the genotype at the B-locus (epistasis). The dominant allele at the B-locus confers resistance to C19, in the right genetic background, i.e. when an individual is double recessive at the A-locus ("aa" genotype). The dominant allele at the C-locus acts epistatically on the A- and B-loci and confers resistance to C1 and C19, regardless of the genotype at the A- and B-loci.

### Supplementary GWAS

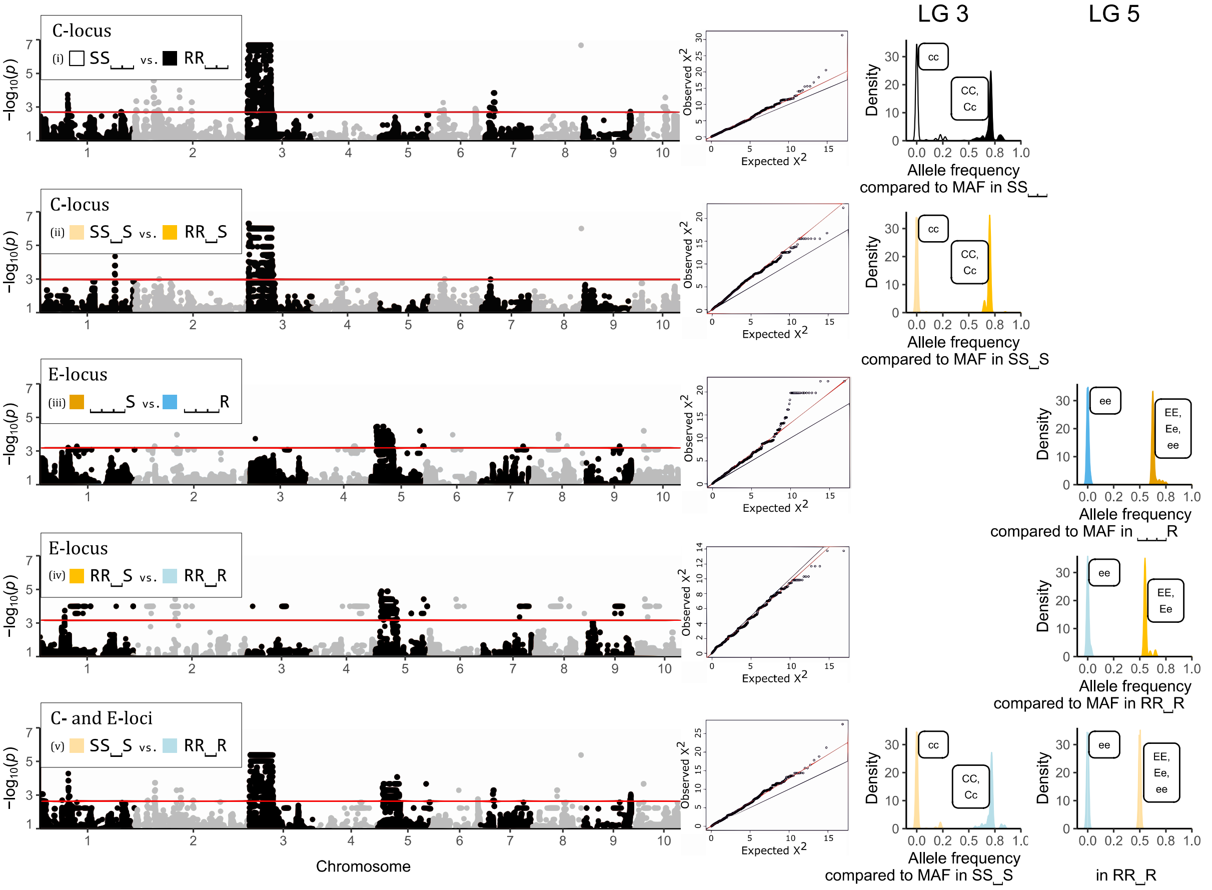

**Figure S3** Comparisons between resistotypes in the Aegelsee *Daphnia magna* population. The resistotype depicts resistance (R) or susceptibility (S) to different *Pasteuria ramosa* isolates in the following order: C1, C19, P15, P20. (i) SS␣␣ vs. RR␣␣; (ii) SS␣S vs. RR␣S; (iii) ␣␣␣S vs. ␣␣␣R; (iv) RR␣S vs. RR␣R, and (v) SS␣S vs. RR␣R. Comparisons (i) and (ii) (variation at C1- and C19-resistotypes) revealed a strong signal on linkage group (LG) 3 corresponding to the C-locus. Comparisons (iii) and (iv) (variation at P20-resistotype) revealed a strong signal on LG 5 corresponding to the E-locus. Comparison (v) (variation at C1- and C19-, and P20-resistotypes) revealed a strong signal on both regions. **Left panel**: Manhattan plots of the resistotype comparisons. The x-axis corresponds to SNP data mapped on the 2.4 *D. magna* reference genome (Routtu et al. 2014), representing only SNPs, not physical distance on the genome. Datapoints with $P_{corrected} < 0.01$ (see Methods) are displayed. **Middle panel**: Quantile-quantile plots of non-corrected p-values excluding SNPs from linkage groups 3 and 5, since these scaffolds displayed an excess of strongly associated markers. **Right panel**: Comparison of allele frequencies between resistotype groups at the C- and the E-loci. Significant SNPs on LG 3 or LG 5 were used (SNPs with $p < P_{lim}/100$, with $P_{lim}$ as defined in the Methods section, equation (1)). For each SNP, the allele with the minor allele frequency (MAF) within resistotype groups that presented only one allele at the C- or the E-locus (all homozygous individuals) was used for comparisons. Hence, the x-axis represents allele frequency of the dominant allele within resistotype groups (considering total allele number, or chromosome number: 2n). Labels attached to peaks describe the inferred possible genotypes at the C- or the E-locus within resistotype groups. In comparisons at the C-locus on LG 3, resistotype groups susceptible to C1 and C19 presented only one allele, i.e. they contained only homozygous recessive individuals at the C-locus (dominant allele frequency of zero). Resistotype groups resistant to C1 and C19 did contain the dominant allele (frequency between 0.5 and 1), showing that resistance is dominant at the C-locus, as the resistant group contains heterozygous individuals. Similarly, in comparisons at the E-locus on LG 5, resistotype groups resistant to P20 do not present the dominant allele (frequency of zero), while resistotype groups susceptible to P20 do, i.e. contain heterozygous individuals (dominant allele frequency between 0.5 and 1). This shows that, in contrast with the C-locus, susceptibility is dominant at the E-locus. Screening individual genomes revealed that some SS␣S individuals (susceptible to C1 and C19, and P20) presented the “ee” genotype at the E-locus (resistance to P20), although susceptibility is dominant at the E-locus. This was not observed in RR␣S individuals (resistant to C1 and C19 but susceptible to P20) (Table S1 below). This observation can be explained by an epistatic relationship linking the C- and the E-loci. This epistasis confers susceptibility to P20 to individuals susceptible to C1 and C19, i.e. presenting the “cc” genotype regardless of the genotype at the E-locus. In contrast with groups containing SS␣S individuals, i.e. comparisons (iii) and (iv), some SS␣S individuals present the “ee” genotype at the E-locus. In these groups, the frequency of the dominant allele can be lower than 0.5.

**Table S1** Genomic description of the E-locus. The region displaying the most markers discriminating between ␣␣␣R and ␣␣␣S encompassed 22 scaffolds and one contig on version 2.4 of the *Daphnia magna* reference genome (Routtu et al. 2014), with a cumulated length of 3101076 bp. The strongest signals of association were found on scaffolds 2167 and 2560. Scaffolds are ordered following the genetic map of (Dukić et al. 2016).

| Scaffold | Length (bp) |
|---|---|
| **scaffold02167** | 358328 |
| contig71262 | 772 |
| scaffold00149 | 66134 |
| scaffold00715 | 344834 |
| scaffold01200 | 81121 |
| scaffold01244 | 6526 |
| scaffold01745 | 65595 |
| scaffold01839 | 55648 |
| scaffold01865 | 165487 |
| scaffold02315 | 259660 |
| scaffold02388 | 172183 |
| scaffold02489 | 70866 |
| **scaffold02560** | 89801 |
| scaffold02708 | 22268 |
| scaffold02913 | 44097 |
| scaffold02970 | 253762 |
| scaffold02993 | 133684 |
| scaffold03031 | 266621 |
| scaffold03154 | 80879 |
| scaffold03356 | 21979 |
| scaffold02031 | 8800 |
| scaffold02207 | 115428 |
| scaffold00872 | 416603 |

**Table S2** We found 485 genes on all associated scaffolds. Scaffolds 2167 and 2560 harbored 82 candidate genes. Some of these genes had annotations similar to genes identified in a previous study of the ABC locus (Bento et al. 2017), with a glucosyltransferase found on scaffold 2167. Three other sugar transferases (galactosyltransferases) were identified on the 22 associated scaffolds, two of them found on scaffold 2560.

| Scaffold | Start | End | Name |
|---|---|---|---|
| scaffold02489 | 49633 | 55225 | udp-galactose:n-acetylgalactosamine-alpha-r beta galactosyltransferase/ARP2_G2977 |
| **scaffold02560** | 1296 | 4007 | udp-galactose:n-acetylgalactosamine-alpha-r beta galactosyltransferase/ARP2_G2977 |
| **scaffold02560** | 4186 | 14123 | udp-galactose:n-acetylgalactosamine-alpha-r beta galactosyltransferase/ARP2_G2977 |
| **scaffold02167** | 334036 | 335960 | glucosyl/glucuronosyl tr_G27 |

**Table S3** List of *D. magna* genotypes used for the association analysis. Genotypes whose name starts with "CH-H" were collected directly from the pond between 2010 and 2014. Names starting with "t" depict animals hatched in the laboratory from resting eggs collected in early 2014, before the natural hatching of the *D. magna* population.

| Clone | Resistotype C1, C19, P15, P20 | Resistotype C1, C19, _ , P20 | C-locus genotype | E-locus genotype |
|---|---|---|---|---|
| CH-H-1769 | RRRR | RR_R | Cc | ee |
| CH-H-2029 | RRRR | RR_R | Cc | ee |
| CH-H-2299 | RRRR | RR_R | CC | ee |
| t1_10.3_2 | RRRR | RR_R | CC | ee |
| t2_14.3_10 | RRRR | RR_R | CC | ee |
| CH-H-1 | RRSR | RR_R | Cc | ee |
| t1_10.3_4 | RRSR | RR_R | Cc | ee |
| t3_10.3_12 | RRSR | RR_R | Cc | unsure |
| t3_14.3_4 | RRSR | RR_R | CC | ee |
| t3_14.3_5 | RRSR | RR_R | Cc | ee |
| t3_31.3_1 | RRSR | RR_R | Cc | ee |
| t4_10.3_11 | RRSR | RR_R | CC | ee |
| t4_10.3_12 | RRSR | RR_R | Cc | ee |
| t4_10.3_2 | RRSR | RR_R | Cc | ee |
| t5_10.3_5 | RRSR | RR_R | CC | ee |
| t5_20.3_1 | RRSR | RR_R | CC | ee |
| t2_17.3_1 | RRSS | RR_S | Cc | Ee |
| t2_17.3_4 | RRSS | RR_S | Cc | Ee |
| t2_17.3_5 | RRSS | RR_S | Cc | EE |
| t3_12.3_1 | RRSS | RR_S | CC | Ee |
| t3_31.3_3 | RRSS | RR_S | CC | EE |
| t4_13.3_10 | RRSS | RR_S | Cc | EE |
| t4_17.3_8 | RRSS | RR_S | Cc | Ee |
| t4_17.3_9 | RRSS | RR_S | CC | Ee |
| t5_12.3_3 | RRSS | RR_S | CC | Ee |
| t5_12.3_4 | RRSS | RR_S | Cc | Ee |
| CH-H-16 | SSSS | SS_S | cc | Ee |
| t2_12.3_10 | SSSS | SS_S | cc | Ee |
| t2_12.3_11 | SSSS | SS_S | cc | ee |
| t2_12.3_12 | SSSS | SS_S | cc | ee |
| t2_14.3_9 | SSSS | SS_S | cc | EE |
| t3_12.3_6 | SSSS | SS_S | cc | ee |
| t3_12.3_8 | SSSS | SS_S | cc | EE |
| t3_13.5_5 | SSSS | SS_S | cc | Ee |
| t3_18.3_3 | SSSS | SS_S | cc | Ee |
| t4_12.3_3 | SSSS | SS_S | cc | EE |
| t4_7.3_4 | SSSS | SS_S | cc | Ee |

### Supplementary Selfing results

#### Selfing in the Aegelsee *Daphnia magna* population: phenotypic and genotypic segregation.

**Tables S4 to S12:** Selfing results for parents with variation at C- and E- loci.
**Tables S13 and S14:** Selfing results for parents with variation at B-, C- and E- loci.
**Table S15:** Detailed results of statistical analyses applied to F1 groups.

**Tables S4 to S12** Expected and observed genotypes and phenotypes of F1 offspring groups resulting from the selfing of F0 parents where variation at the C- and E-loci was observed (see Supplementary selfing methods Table S21 for clones' details). Observed resistance genotype for parents and offspring was assessed using markers DMPR1 and DMPR3 for the C- and the E-loci, respectively (see results). Resistance is dominant at the C-locus (resistance to C1/C19) whereas resistance is recessive at the E-locus (resistance to P20). In addition, an epistatic relationship linking both loci confers susceptibility to P20 to an individual that shows susceptibility to C1 and C19, disregarding the individual genotype at the E-locus (see results section Fig. 5). Text in red denotes genotypes and their corresponding phenotypes where the epistatic relationship between the C and the E-loci is expected to be revealed in the phenotype. The epistatic relationship "confers" to "cc--" individuals a susceptibility to P20, whatever their genotype at the E-locus. Hence the epistasis can only be observed phenotypically in "ccee" offspring. If the epistatic relationship is not present in this case, the observed phenotype would be SS␣R, whereas if the epistatic relationship is present, the observed phenotype would be SS␣S. **A.** Expected Punnett square for the selfed genotype according to the genetic model. **B.** Expected vs. observed genotypes and phenotypes of selfed F1 offspring. Differences in offspring number between the genotype and the phenotype correspond to instances where either genetic markers analysis did not work, or attachment test was not conducted because the *D. magna* clone got extinct. Expected genotypes and phenotypes were calculated from the Punnett square and using the R package "peas" (Supplementary peas Doc. S1). Genotypes are ordered in upper-case then lower-case fashion. We compared expected vs. observed genotype and resistotype segregation separately in the F1 groups using the Cochran–Mantel–Haenszel (C-M-H) test for repeated tests of independence. When there was only one repeat (Table S7), we used the Fisher test. In cases where there was only one category of expected and observed genotype or phenotype, no test was run (Tables S4, S6, S10, S11 (resistotype only) and S12). In these cases, expectation and observation show a perfect match. Following each C-M-H test, assumption of homogeneity of the odds ratio across repeats was confirmed using a Breslow-Day test (R package DescTools: (Signorell et al. 2018)). However, this test can only be operated in 2x2 tables (phenotype only Tables S5, S7, S8). We then ran a Fisher test of independence for each comparison (expected vs. observed for each repeat, Bonferroni corrected) to detect possible significant differences in opposite directions across repeats, which would result in a non-significant C-M-H test. We did not detect such differences in direction (see Table S15 below for all detailed results of statistical analyses). Tests were run on counts, although we present here segregation of offspring as proportions.

**Table S4** Selfed genotype "CCEE". F0 parents: **a**: t3_12.3_1i_12, **b**: t3_12.3_1i_21.

**A**

| **RR␣S CCEE** | **CE** |
|---|---|
| **CE** | RR␣S |

**B**

| **Markers genotype** | Fraction: Expected | Fraction: Observed, Repeat **a** n = 43 | Fraction: Observed, Repeat **b** n = 37 | C-M-H test on counts | **Resistotype** | Fraction: Expected | Fraction: Observed, Repeat **a** n = 43 | Fraction: Observed, Repeat **b** n = 37 | C-M-H test on counts |
|---|---|---|---|---|---|---|---|---|---|
| CCEE | 1 | 1 | 1 | NA | RR␣S | 1 | 1 | 1 | NA |

**Table S5** Selfed genotype "CCEe". F0 parents: **a**: CH-2015-36, **b**: t3_12.3_1, **c**: CH-H-2015-49. Repeat "c" was not tested genotypically as the E-locus marker appeared not to be linked to the E-locus.

**A**

| RR␣S CCEe | CE | Ce |
|---|---|---|
| CE | RR␣S | RR␣S |
| Ce | RR␣S | RR␣R |

**B**

| Markers genotype | Fraction: Expected | Fraction: Observed, Repeat **a** n = 89 | Fraction: Observed, Repeat **b** n = 31 | Fraction: Observed, Repeat **c** n = 81 | C-M-H test on counts (a+b) | Resistotype | Fraction: Expected | Fraction: Observed, Repeat **a** n = 89 | Fraction: Observed, Repeat **b** n = 31 | Fraction: Observed, Repeat **c** n = 79 | C-M-H test on counts |
|---|---|---|---|---|---|---|---|---|---|---|---|
| CCEE | 0.25 | 0.31 | 0.19 | 1 | | RR␣S | 0.75 | 0.79 | 0.74 | 0.84 | |
| CCEe | 0.50 | 0.46 | 0.55 | 0 | $M^2 = 0.35$, $df = 2$, $p = 0.84$ | | | | | | $X^2 = 0.85$, $df = 1$, $p = 0.36$ |
| CCee | 0.25 | 0.22 | 0.26 | 0 | | RR␣R | 0.25 | 0.21 | 0.26 | 0.16 | |

**Table S6** Selfed genotype "CCee". F0 parents: **a**: CH-H-434-inb2-1, **b**: t1_10.3_2, **c**: CH-H-2015-16, **d**: CH-H-2016-b-70.

**A**

| RR␣R CCee | Ce |
|---|---|
| Ce | RR␣R |

**B**

| Markers genotype | Fraction: Expected | Fraction: Observed, Repeat **a** n = 39 | Fraction: Observed, Repeat **b** n = 42 | Fraction: Observed, Repeat **c** n= 66 | Fraction: Observed, Repeat **d** n = 84 | C-M-H test on counts | Resistotype | Fraction: Expected | Fraction: Observed, Repeat **a** n = 39 | Fraction: Observed, Repeat **b** n = 42 | Fraction: Observed, Repeat **c** n = 70 | Fraction: Observed, Repeat **d** n = 79 | C-M-H test on counts |
|---|---|---|---|---|---|---|---|---|---|---|---|---|---|
| CCee | 1 | 1 | 1 | 1 | 1 | NA | RR␣R | 1 | 1 | 1 | 1 | 1 | NA |

**Table S7** Selfed genotype "CcEE". F0 parent: **a**: t2_17.3_4i_12.

**A**

| RR␣S CcEE | CE | cE |
|---|---|---|
| CE | RR␣R | RR␣R |
| cE | RR␣R | SS␣S |

**B**

| Markers genotype | Fraction: Expected | Fraction: Observed, Repeat **a** n = 19 | Fisher test on counts | Resistotype | Fraction: Expected | Fraction: Observed, Repeat **a** n = 19 | Fisher test on counts |
|---|---|---|---|---|---|---|---|
| CCEE | 0.25 | 0.32 | | RR␣S | 0.75 | 0.74 | |
| CcEE | 0.50 | 0.42 | $p = 0.92$ | | | | $p = 1$ |
| ccEE | 0.25 | 0.26 | | SS␣S | 0.25 | 0.26 | |

**Table S8** Selfed genotype "CcEe". F0 parents: **a**: t3_14.3_1, **b**: t2_17.3_4, **c**: t2_17.3_1, **d**: CH-H-2015-59. Repeat "d" was not tested genotypically as the E-locus marker appeared not to be linked to the E-locus. Text in grey corresponds to non-matching marker genotypes with their observed resistotypes according to the genetic model. Those non-matching observations are included above in the observed proportions of genotypes.

**A**

| RR_S CcEe | CE | Ce | cE | ce |
|---|---|---|---|---|
| CE | RR_S | RR_S | RR_S | RR_S |
| Ce | RR_S | RR_R | RR_S | RR_R |
| cE | RR_S | RR_S | SS_S | SS_S |
| ce | RR_S | RR_R | SS_S | SS_S |

**B**

| Markers genotype | Fraction: Expected | Fraction: Observed, Repeat **a** n = 47 | **b** n = 32 | **c** n= 80 | **d** n = 61 | C-M-H test on counts (a+b+c) | Resistotype | Fraction: Expected | Fraction: Observed, Repeat **a** n = 48 | **b** n = 34 | **c** n = 64 | **d** n = 65 | C-M-H test on counts |
|---|---|---|---|---|---|---|---|---|---|---|---|---|---|
| CCEE | 0.06 | 0 | 0.16 | 0.05 | 0.20 | $M^2$ = 6.79, *df* = 8, *p* = 0.56 | RR_S | 0.56 | 0.47 | 0.56 | 0.65 | 0.51 | $M^2$ = 4.61, *df* = 2, *p* = 0.10 |
| CCEe | 0.13 | 0 | 0.19 | 0.12 | 0 | | | | | | | | |
| CcEE | 0.13 | 0.13 | 0.06 | 0.18 | 0.57 | | | | | | | | |
| CcEe | 0.25 | 0.40 | 0.25 | 0.28 | 0 | | | | | | | | |
| ccEE | 0.06 | 0.09 | 0 | 0.03 | 0.23 | | SS_S | 0.25 | 0.30 | 0.06 | 0.13 | 0.21 | |
| ccEe | 0.13 | 0.11 | 0.03 | 0.03 | 0 | | | | | | | | |
| ccee | 0.06 | 0.11 | 0.03 | 0.03 | 0 | | | | | | | | |
| CCee | 0.06 | 0 | 0.09 | 0.17 | 0 | | RR_R | 0.19 | 0.23 | 0.38 | 0.22 | 0.28 | |
| Ccee | 0.13 | 0.17 | 0.19 | 0.10 | 0 | | | | | | | | |
| CCEE | | | | | 0.11 | | RR_R | | | | | | |
| CCEe | | | 0.06 | | | | RR_R | | | | | | |
| CcEE | | 0.02 | 0.03 | | 0.16 | | RR_R | | | | | | |
| CcEe | | 0.06 | 0.06 | | | | RR_R | | | | | | |
| CCee | | | | 0.05 | | | RR_S | | | | | | |
| Ccee | | 0.02 | 0.06 | | | | RR_S | | | | | | |
| CcEE | | | | 0.02 | 0.02 | | SS_S | | | | | | |
| CcEe | | | | 0.02 | | | SS_S | | | | | | |
| ccEE | | | | | 0.03 | | RR_S | | | | | | |

**Table S9** Selfed genotype "Ccee". F0 parents: **a**: t1_10.3_4, **b**: t5_10.3_3, **c**: CH-H-434-inb2-2.

**A**

| RR_R Ccee | Ce | ce |
|---|---|---|
| Ce | RR_R | RR_R |
| ce | RR_R | SS_S |

**B**

| Markers genotype | Fraction: Expected | Fraction: Observed, Repeat **a** n = 36 | **b** n = 49 | **c** n= 21 | C-M-H test on counts | Resistotype | Fraction: Expected | Fraction: Observed, Repeat **a** n = 36 | **b** n = 49 | **c** n= 22 | C-M-H test on counts |
|---|---|---|---|---|---|---|---|---|---|---|---|
| CCee | 0.25 | 0.19 | 0.16 | 0.14 | $M^2$ = 2.16, *df* = 2, *p* = 0.34 | RR_R | 0.75 | 0.75 | 0.80 | 0.62 | $X^2$ = 0.0062, *df* = 1, *p* = 0.94 |
| Ccee | 0.50 | 0.56 | 0.63 | 0.48 | | | | | | | |
| ccee | 0.25 | 0.25 | 0.20 | 0.38 | | SS_S | 0.25 | 0.25 | 0.20 | 0.38 | |

**Table S10** Selfed genotype "ccEE". F0 parent: **a**: CH-H-2015-97.

**A**

| SS␣S ccEE | cE |
|---|---|
| cE | SS␣S |

**B**

| Markers genotype | Fraction | | Fisher test on counts | Resistotype | Fraction | | Fisher test on counts |
|---|---|---|---|---|---|---|---|
| | Expected | Observed<br>Repeat<br>**a**<br>n = 89 | | | Expected | Observed<br>Repeat<br>**a**<br>n = 87 | |
| ccEE | 1 | 1 | NA | SS␣S | 1 | 1 | NA |

**Table S11** Selfed genotype "ccEe". F0 parents: **a**: CH-H-2015-113, **b**: t4_10.3_16.

**A**

| SS␣S ccEe | cE | ce |
|---|---|---|
| cE | SS␣S | SS␣S |
| ce | SS␣S | SS␣S |

**B**

| Markers genotype | Fraction | | | C-M-H test on counts | Resistotype | Fraction | | | C-M-H test on counts |
|---|---|---|---|---|---|---|---|---|---|
| | Expected | Observed<br>Repeat<br>**a**<br>n = 82 | Observed<br>Repeat<br>**b**<br>n = 63 | | | Expected | Observed<br>Repeat<br>**a**<br>n = 84 | Observed<br>Repeat<br>**b**<br>n = 65 | |
| ccEE | 0.25 | 0.27 | 0.15 | $M^2 = 0.58$, $df = 2$, $p = 0.75$ | SS␣S | 1 | 1 | 1 | NA |
| ccEe | 0.50 | 0.55 | 0.44 | | | | | | |
| ccee | 0.25 | 0.18 | 0.41 | | | | | | |

**Table S12** Selfed genotype "ccee". F0 parents: **a**: t4_13.3_2, **b**: CH-H-2015-86.

**A**

| SS␣S ccee | ce |
|---|---|
| ce | SS␣S |

**B**

| Markers genotype | Fraction | | | C-M-H test on counts | Resistotype | Fraction | | | C-M-H test on counts |
|---|---|---|---|---|---|---|---|---|---|
| | Expected | Observed<br>Repeat<br>**a**<br>n = 76 | Observed<br>Repeat<br>**b**<br>n = 36 | | | Expected | Observed<br>Repeat<br>**a**<br>n = 74 | Observed<br>Repeat<br>**b**<br>n = 35 | |
| ccee | 1 | 1 | 1 | NA | SS␣S | 1 | 1 | 1 | NA |

**Tables S13 and S14** Expected and observed phenotypes of F1 offspring resulting from the selfing of F0 parents where variation at the B- and E-loci was observed (see Supplementary selfing methods Table S21 for clones' details). Individuals bearing the dominant allele at the B-locus display a SR␣␣ resistotype underlined by a "--B-cc" genotype. In the Swisspond population we assume the recessive allele is fixed at the A-locus and the dominant allele at the B-locus is rare (see results section). Individuals with a SR␣␣ resistotype are thus expected to possess an "aaB-cc" genotype in our study population. As genetic markers were not designed to discriminate variation at the B-locus and consequently do not show linkage with expected genotypes, we present only phenotype segregation. We deduced the resistance genotypes of the present F0 parents based on their resistotype and on the segregation patterns of their F1 offspring. In these F1 offspring, we observed SR␣R individuals, supposedly "aaB-ccee", which indicates that the epistatic relationship described before between the C- and the E-loci (results section Fig. 5) should also include the B-locus. If the epistasis was only between the C- and the E-locus, SR␣␣ individuals would necessarily be SR␣S. However, we do find SR␣R individuals among F1 offspring of SR␣S parents, which indicates that the "bbcc--" genotype induces susceptibility to P20, regardless of the genotype at the E-locus. As SR␣␣ individuals are rare in the population, we base this hypothesis solely on the selfed offspring presented below. As the recessive allele at the A-locus is fixed in the population, we cannot infer a possible role of the A-locus in this epistatic relationship. Text in red denotes phenotypes and genotypes where this epistatic relationship between the B/C- and the E-loci is expected to be revealed in the phenotype. The epistatic relationship "confers" to "bbcc--" individuals a susceptibility to P20, whatever their genotype at the E-locus. Hence the epistasis can only be observed phenotypically in "bbccee" offspring. If the epistatic relationship was not present in this case, the observed phenotype would be SS␣R, whereas if the epistatic relationship is present, the observed phenotype is SS␣S. This genetic model is presented in Fig. 5 in the results section. **A.** Expected Punnett square for the selfed genotype according to the genetic model including the B-locus. **B.** Expected and observed phenotypes of selfed F1 offspring. Expected phenotypes were calculated from the Punnett square and using the R package "peas" (Supplementary peas Doc. S2). Genotypes are ordered in upper-case then lower-case fashion. We compared expected vs. observed resistotype segregation in the F1 groups using the Fisher test of independence (see Table S15 below for all detailed results of statistical analyses). Tests were run on counts, although we present here segregation of offspring as proportions.

**Table S13** Selfed genotype "BbccEE". F0 parent: **a**: t0_9.3_7.

**A**

| SR␣S BbccEE | BcE | bcE |
|---|---|---|
| BcE | SR␣S | SR␣S |
| bcE | SR␣S | SS␣S |

**B**

| Markers genotype | Fraction: Expected | Fraction: Observed, Repeat **a** n = 35 | Fisher test on counts | Resistotype | Fraction: Expected | Fraction: Observed, Repeat **a** n = 37 | Fisher test on counts |
|---|---|---|---|---|---|---|---|
| BBccEE | 0.25 | NA | | SR␣S | 0.75 | 0.76 | |
| BbccEE | 0.50 | NA | NA | SS␣S | 0.25 | 0.22 | $p = 1$ |
| bbccEE | 0.25 | NA | | SR␣R | 0.00 | 0.03 | |

**Table S14** Selfed genotype “BbccEe”. F0 parent: **a**: t0_28.2_43.

**A**

| SR␣S BbccEe | BcE | Bce | bcE | bce |
|---|---|---|---|---|
| BcE | SR␣S | SR␣S | SR␣S | SR␣S |
| Bce | SR␣S | SR␣R | SR␣S | SR␣R |
| bcE | SR␣S | SR␣S | SS␣S | SS␣S |
| bce | SR␣S | SR␣R | SS␣S | SS␣S |

**B**

| Markers genotype | Fraction: Expected | Fraction: Observed, Repeat a (n = 36) | Fisher test on counts | Resistotype | Fraction: Expected | Fraction: Observed, Repeat a (n = 38) | Fisher test on counts |
|---|---|---|---|---|---|---|---|
| BBccEE | 0.06 | NA | | SR␣S | 0.56 | 0.55 | |
| BBccEe | 0.13 | NA | | | | | |
| BbccEE | 0.13 | NA | | | | | |
| BbccEe | 0.25 | NA | | | | | |
| bbccEE | 0.06 | NA | NA | SS␣S | 0.25 | 0.32 | $p = 0.58$ |
| bbccEe | 0.13 | NA | | | | | |
| bbccee | 0.06 | NA | | | | | |
| BBccee | 0.06 | NA | | SR␣R | 0.19 | 0.13 | |
| Bbccee | 0.13 | NA | | | | | |

**Table S15** Detailed results of statistical analyses applied to F1 offspring groups. Analyses are described in tables’ legends above.

#### Phenotype

| | CMH test | | | | BD test | | | Fisher test | | | | | | | |
|---|---|---|---|---|---|---|---|---|---|---|---|---|---|---|---|
| | | | | | | | | repeat a | | repeat b | | repeat c | | repeat d | |
| | *M2* | *df* | *p* | | *X²* | *df* | *p* | *p* | *p + Bonf.* | *p* | *p + Bonf.* | *p* | *p + Bonf.* | *p* | *p + Bonf.* |
| table 1 | NA | NA | NA | NA | NA | NA | NA | NA | NA | NA | NA | NA | NA | NA | NA |
| table 2 | X² = 0.85 | 1 | 0.36 | CI = (0.49, 1.25), estimate = 0.78, null value = 1 | 1 | 2 | 1 | 1 | 1 | 1 | 1 | 0 | 0.72 | NA | NA |
| table 3 | NA | NA | NA | NA | NA | NA | NA | NA | NA | NA | NA | NA | NA | NA | NA |
| table 4 | NA | NA | NA | NA | NA | NA | NA | 1 | NA | NA | NA | NA | NA | NA | NA |
| table 5 | 4.61 | 2 | 0.1 | NA | NA | NA | NA | 1 | 1 | 0 | 0.27 | 0 | 1 | 0 | 1 |
| table 6 | X² = 0.0062 | 1 | 0.94 | CI = (0.55, 1.9), estimate = 1.03, null value = 1 | 1 | 2 | 1 | 1 | 1 | 1 | 1 | 1 | 1 | NA | NA |
| table 7 | NA | NA | NA | NA | NA | NA | NA | NA | NA | NA | NA | NA | NA | NA | NA |
| table 8 | NA | NA | NA | NA | NA | NA | NA | NA | NA | NA | NA | NA | NA | NA | NA |
| table 9 | NA | NA | NA | NA | NA | NA | NA | NA | NA | NA | NA | NA | NA | NA | NA |
| table 10 | NA | NA | NA | NA | NA | NA | NA | 1 | NA | NA | NA | NA | NA | NA | NA |
| table 11 | NA | NA | NA | NA | NA | NA | NA | 1 | NA | NA | NA | NA | NA | NA | NA |

#### Genotype

| | CMH test | | | Fisher test | | | | | |
|---|---|---|---|---|---|---|---|---|---|
| | | | | repeat a | | repeat b | | repeat c | |
| | *M²* | *df* | *p* | *p* | *p + Bonf.* | *p* | *p + Bonf.* | *p* | *p + Bonf.* |
| table 1 | NA | NA | NA | NA | NA | NA | NA | NA | NA |
| table 2 | 0.35 | 2 | 0.84 | 1 | 1 | 1 | 1 | NA | NA |
| table 3 | NA | NA | NA | NA | NA | NA | NA | NA | NA |
| table 4 | NA | NA | NA | 1 | NA | NA | NA | NA | NA |
| table 5 | 6.79 | 8 | 0.56 | 0 | 0.19 | 1 | 1 | 0 | 1 |
| table 6 | 2.16 | 2 | 0.34 | 1 | 1 | 0 | 1 | 1 | 1 |
| table 7 | NA | NA | NA | NA | NA | NA | NA | NA | NA |
| table 8 | 0.58 | 2 | 0.75 | 1 | 1 | 0 | 0.25 | NA | NA |
| table 9 | NA | NA | NA | NA | NA | NA | NA | NA | NA |
| table 10 | NA | NA | NA | NA | NA | NA | NA | NA | NA |
| table 11 | NA | NA | NA | NA | NA | NA | NA | NA | NA |

### Supplementary Markers linkage

#### Multi-marker genotypes and their corresponding resistance genotypes and phenotypes: expected vs. observed.

**Table S16** Multi-marker genotypes and their expected corresponding resistance genotype at the C and E-loci and resistotypes (resistance phenotypes) of *Daphnia magna* in the Swisspond population, assuming perfect linkage between markers and resistance loci. All possible allele combinations for the four markers DMPR1, 2, 3 and 4 (DMPR: *Daphnia magna* - *Pasteuria ramosa*). DMPR1 and 2 are physically linked to the C-locus while DMPR3 and 4 are physically linked to the E-locus. Zeros (0) and ones (1) correspond to absence and presence of an allele, respectively. The size of an amplicon containing an allele is denoted next to the name of the allele (see Supplementary GWAS methods for details on the markers). For example, DMPR1_R1_89 corresponds to the amplicon of 89 bp yielded by the PCR for the first R allele of the DMPR1 marker. Inferred resistotypes in red denote resistotypes where the epistatic interaction between the C and the E-loci is detectable. In order to show the logic of the different possible combinations, lines with zeros and ones identical to the line directly above them are shown in grey.

| Alleles associated to the C-locus | | | | | Alleles associated to the E-locus | | | | | | | |
|---|---|---|---|---|---|---|---|---|---|---|---|---|
| DMPR1_R1_89 | DMPR1_R2_113 | DMPR1_S_118 | DMPR2_S_176 | DMPR2_R_184 | DMPR3_S_206 | DMPR3_R1_211 | DMPR3_R2_212 | DMPR4_R_128 | DMPR4_S_136 | Multi-marker genotype | Inferred resistance genotype | Inferred resistotype |
| **0** | **0** | **1** | **1** | **0** | **0** | **0** | **1** | **1** | **0** | 001_10_001_10 | ccee | SS_S |
| 0 | 0 | 1 | 1 | 0 | **0** | **1** | **0** | 1 | 0 | 001_10_010_10 | ccee | SS_S |
| 0 | 0 | 1 | 1 | 0 | **0** | **1** | **1** | 1 | 0 | 001_10_011_10 | ccee | SS_S |
| 0 | 0 | 1 | 1 | 0 | **1** | **0** | **0** | **0** | **1** | 001_10_100_01 | ccEE | SS_S |
| 0 | 0 | 1 | 1 | 0 | **1** | **0** | **1** | **1** | **1** | 001_10_101_11 | ccEe | SS_S |
| 0 | 0 | 1 | 1 | 0 | **1** | **1** | **0** | 1 | 1 | 001_10_110_11 | ccEe | SS_S |
| **0** | **1** | **0** | **0** | **1** | **0** | **0** | **1** | **1** | **0** | 010_01_001_10 | CCee | RR_R |
| 0 | 1 | 0 | 0 | 1 | **0** | **1** | **0** | 1 | 0 | 010_01_010_10 | CCee | RR_R |
| 0 | 1 | 0 | 0 | 1 | **0** | **1** | **1** | 1 | 0 | 010_01_011_10 | CCee | RR_R |
| 0 | 1 | 0 | 0 | 1 | **1** | **0** | **0** | **0** | **1** | 010_01_100_01 | CCEE | RR_S |
| 0 | 1 | 0 | 0 | 1 | **1** | **0** | **1** | **1** | **1** | 010_01_101_11 | CCEe | RR_S |
| 0 | 1 | 0 | 0 | 1 | **1** | **1** | **0** | 1 | 1 | 010_01_110_11 | CCEe | RR_S |
| **0** | **1** | **1** | **1** | **1** | **0** | **0** | **1** | **1** | **0** | 011_11_001_10 | Ccee | RR_R |
| 0 | 1 | 1 | 1 | 1 | **0** | **1** | **0** | 1 | 0 | 011_11_010_10 | Ccee | RR_R |
| 0 | 1 | 1 | 1 | 1 | **0** | **1** | **1** | 1 | 0 | 011_11_011_10 | Ccee | RR_R |
| 0 | 1 | 1 | 1 | 1 | **1** | **0** | **0** | **0** | **1** | 011_11_100_01 | CcEE | RR_S |
| 0 | 1 | 1 | 1 | 1 | **1** | **0** | **1** | **1** | **1** | 011_11_101_11 | CcEe | RR_S |
| 0 | 1 | 1 | 1 | 1 | **1** | **1** | **0** | 1 | 1 | 011_11_110_11 | CcEe | RR_S |
| **1** | **0** | **0** | **0** | **1** | **0** | **0** | **1** | **1** | **0** | 100_01_001_10 | CCee | RR_R |
| 1 | 0 | 0 | 0 | 1 | **0** | **1** | **0** | 1 | 0 | 100_01_010_10 | CCee | RR_R |
| 1 | 0 | 0 | 0 | 1 | **0** | **1** | **1** | 1 | 0 | 100_01_011_10 | CCee | RR_R |
| 1 | 0 | 0 | 0 | 1 | **1** | **0** | **0** | **0** | **1** | 100_01_100_01 | CCEE | RR_S |
| 1 | 0 | 0 | 0 | 1 | **1** | **0** | **1** | **1** | **1** | 100_01_101_11 | CCEe | RR_S |
| 1 | 0 | 0 | 0 | 1 | **1** | **1** | **0** | 1 | 1 | 100_01_110_11 | CCEe | RR_S |
| **1** | **0** | **1** | **1** | **1** | **0** | **0** | **1** | **1** | **0** | 101_11_001_10 | Ccee | RR_R |
| 1 | 0 | 1 | 1 | 1 | **0** | **1** | **0** | 1 | 0 | 101_11_010_10 | Ccee | RR_R |
| 1 | 0 | 1 | 1 | 1 | **0** | **1** | **1** | 1 | 0 | 101_11_011_10 | Ccee | RR_R |
| 1 | 0 | 1 | 1 | 1 | **1** | **0** | **0** | **0** | **1** | 101_11_100_01 | CcEE | RR_S |
| 1 | 0 | 1 | 1 | 1 | **1** | **0** | **1** | **1** | **1** | 101_11_101_11 | CcEe | RR_S |
| 1 | 0 | 1 | 1 | 1 | **1** | **1** | **0** | 1 | 1 | 101_11_110_11 | CcEe | RR_S |
| **1** | **1** | **0** | **0** | **1** | **0** | **0** | **1** | **1** | **0** | 110_01_001_10 | CCee | RR_R |
| 1 | 1 | 0 | 0 | 1 | **0** | **1** | **0** | 1 | 0 | 110_01_010_10 | CCee | RR_R |
| 1 | 1 | 0 | 0 | 1 | **0** | **1** | **1** | 1 | 0 | 110_01_011_10 | CCee | RR_R |
| 1 | 1 | 0 | 0 | 1 | **1** | **0** | **0** | **0** | **1** | 110_01_100_01 | CCEE | RR_S |
| 1 | 1 | 0 | 0 | 1 | **1** | **0** | **1** | **1** | **1** | 110_01_101_11 | CCEe | RR_S |
| 1 | 1 | 0 | 0 | 1 | **1** | **1** | **0** | 1 | 1 | 110_01_110_11 | CCEe | RR_S |

**Table S17** Observed allele combinations for markers DMPR1 and 2 and their corresponding inferred resistance genotype, expected and observed resistotypes. DMPR1 and 2 are linked to the C-locus. Details about the markers and their expected signatures from the genetic model of resistance are given in Supplementary GWAS methods. Text in grey represents occurrences where the measured and expected resistotypes do not match. For each genotype, the expected count of resistotypes is calculated as the total count of observed resistotypes. The panel of clones used represents 24 selfed offspring groups (F1) ranging from 19 to 89 individual clones (Supplementary selfing methods), as well as three random samples from the Swisspond from 2014 to 2016 (22 to 108 clones), either hatched from resting eggs of directly sampled from the pond. Differences in total sample size correspond to occurrences where a marker did not amplify or when the marker signature could not be clearly identified. The initial total sample size of phenotyped clones is n = 1550. Individuals with a "SR_ _" resistotype were excluded from the analysis as they were not used in the GWAS from which the markers were designed. Better linkage is observed between the C-locus and DMPR1 than between the C-locus and DMPR2.

| DMPR1_R1_89 | DMPR1_R2_113 | DMPR1_S_118 | DMPR1 genotype | Inferred resistance genotype | Inferred resistotype | Measured resistotype | Count (n = 1442) | Expected count from inferred resistotypes | Frequency (n = 1442) | Expected frequency from inferred resistotypes |
|---|---|---|---|---|---|---|---|---|---|---|
| **0** | **0** | **1** | 001 | cc | SS_ _ | SS_ _ | 488 | 490 | 0.338 | 0.340 |
| | | | | | | *RR_ _* | *2* | *0* | *0.001* | *0* |
| **0** | **1** | **0** | 010 | CC | RR_ _ | RR_ _ | 515 | 515 | 0.357 | 0.357 |
| **0** | **1** | **1** | 011 | Cc | RR_ _ | RR_ _ | 251 | 254 | 0.174 | 0.176 |
| | | | | | | *SS_ _* | *3* | *0* | *0.002* | *0* |
| **1** | **0** | **0** | 100 | CC | RR_ _ | RR_ _ | 76 | 76 | 0.053 | 0.053 |
| **1** | **0** | **1** | 101 | Cc | RR_ _ | RR_ _ | 67 | 68 | 0.046 | 0.047 |
| | | | | | | *SS_ _* | *1* | *0* | *0.001* | *0* |
| **1** | **1** | **0** | 110 | CC | RR_ _ | RR_ _ | 39 | 39 | 0.027 | 0.027 |
| | | | | | | | Fisher's exact test | | | |
| | | | | | | | *p* = 0.77 | | | |

| DMPR2_S_176 | DMPR2_R_184 | | DMPR2 genotype | Inferred resistance genotype | Inferred resistotype | Measured resistotype | Count (n = 1434) | Expected count from inferred resistotypes | Frequency (n = 1434) | Expected frequency from inferred resistotypes |
|---|---|---|---|---|---|---|---|---|---|---|
| **0** | **1** | | 01 | CC | RR_ _ | RR_ _ | 548 | 549 | 0.382 | 0.383 |
| | | | | | | *SS_ _* | *1* | *0* | *0.001* | *0* |
| **1** | **0** | | 10 | cc | SS_ _ | SS_ _ | 480 | 599 | 0.335 | 0.418 |
| | | | | | | *RR_ _* | *119* | *0* | *0.083* | *0* |
| **1** | **1** | | 11 | Cc | RR_ _ | RR_ _ | 283 | 286 | 0.197 | 0.199 |
| | | | | | | *SS_ _* | *3* | *0* | *0.002* | *0* |
| | | | | | | | Fisher's exact test | | | |
| | | | | | | | $p = 1.25.10^{-37}$ | | | |

**Table S18** Observed allele combinations for markers DMPR3 and 4 and their corresponding inferred resistance genotype, expected and observed resistotypes. DMPR1 and 2 are linked to the E-locus. Details about the markers and their expected signatures from the genetic model of resistance are given in Supplementary GWAS methods. Within the groups of "ee" genotypes, we separately present the "SS__" C1/C19 resistotype as we expect in that case the P20 resistotype to be "___S", due to the epistatic relationship between the C and the E-locus. Text in grey represents occurrences where the measured and expected resistotypes do not match. Text in red denotes resistotypes where the epistatic interaction between the C and the E-loci is detectable. For each genotype, the expected count of resistotypes is calculated as the total count of observed resistotypes. The panel of clones used represents 24 selfed offspring groups (F1) ranging from 12 to 84 individual clones (Supplementary selfing methods), as well as three random samples from the Swisspond from 2014 to 2016 (22 to 108 clones), either hatched from resting eggs of directly sampled in the pond. Differences in total sample size correspond to occurrences where a marker did not amplify or when the marker signature could not be clearly identified. The initial total sample size of phenotyped clones is n = 1550. Individuals with a "SR__" resistotype were excluded from the analysis as they were not used in the GWAS from which the markers were designed. Better linkage is observed between the E-locus and DMPR3 than between the E-locus and DMPR4. Note that marker DMPR4 did not amplify as well as the other markers (n = 911). This marker showed better amplification when put alone in a PCR reaction however we decided not to use it further as it showed poor linkage to the E-locus.

| DMPR3_S_206 | DMPR3_R1_211 | DMPR3_R2_212 | DMPR3 genotype | Inferred resistance genotype | Measured resistotype to C1/C19 | Inferred resistotype to P20 | Measured resistotype to P20 | Count (n = 1438) | Expected count from inferred resistotypes | Frequency (n = 1438) | Expected frequency from inferred resistotypes |
|---|---|---|---|---|---|---|---|---|---|---|---|
| **0** | **1** | **0** | 010 | ee | RR__ | ___R | ___R | 468 | 481 | 0.325 | 0.334 |
| | | | | | | | *___S* | *13* | *0* | *0.009* | *0* |
| | | | | | SS__ | ___S | ___S | 227 | 227 | 0.158 | 0.158 |
| | | | | | | | *___R* | | | | |
| **0** | **1** | **1** | 011 | ee | RR__ | ___R | ___R | 3 | 3 | 0.002 | 0.002 |
| **1** | **0** | **0** | 100 | EE | XX__ | ___S | ___S | 417 | 450 | 0.290 | 0.313 |
| | | | | | | | *___R* | *33* | *0* | *0.023* | *0* |
| **1** | **0** | **1** | 101 | Ee | XX__ | ___S | ___S | 1 | 1 | 0.001 | 0.001 |
| **1** | **1** | **0** | 110 | Ee | XX__ | ___S | ___S | 247 | 276 | 0.172 | 0.192 |
| | | | | | | | *___R* | *29* | *0* | *0.020* | *0* |
| | | | | | | | | Fisher's exact test $p = 5.34.10^{-19}$ | | | |

| DMPR4_R_128 | DMPR4_S_136 | | DMPR4 genotype | Inferred resistance genotype | Measured resistotype to C1/C19 | Inferred resistotype to P20 | Measured resistotype to P20 | Count (n = 911) | Expected count from inferred resistotypes | Frequency (n = 911) | Expected frequency from inferred resistotypes |
|---|---|---|---|---|---|---|---|---|---|---|---|
| **0** | **1** | | 01 | EE | XX__ | ___S | ___S | 357 | 374 | 0.392 | 0.411 |
| | | | | | | | *___R* | *17* | *0* | *0.019* | *0* |
| **1** | **0** | | 10 | ee | RR__ | ___R | ___R | 18 | 19 | 0.020 | 0.021 |
| | | | | | | | *___S* | *1* | *0* | *0.001* | *0* |
| | | | | | SS__ | ___S | ___S | 6 | 6 | 0.007 | 0.007 |
| **1** | **1** | | 11 | Ee | XX__ | ___S | ___S | 251 | 512 | 0.276 | 0.562 |
| | | | | | | | *___R* | *261* | *0* | *0.286* | *0* |
| | | | | | | | | Fisher's exact test $p = 1.00.10^{-100}$ | | | |

#### Supplementary Allele frequency

Natural polymorphism at the C- and E-loci in the Aegelsee *Daphnia magna* population. This study explores the evolution of resistance genotype and allele frequency over the active season of *Daphnia magna* in the Aegelsee (Results section Fig. 6), using (i) observed longitudinal resistotype (resistance phenotype) data (Results section Fig. 1), (ii) genotype and allele polymorphism at resistance loci (the C- and E-loci) in a spring sample of the host population and (iii) the genetic model of resistance yielded in the present study (Results section Fig. 5). **Table S19** presents the genetic model of resistance, considering the C- and the E-loci, in the form of genotype-phenotype pairs. **Figure S4** presents the observed distribution of genotype-phenotype pairs in the host sample and **Table S20** presents the observed distribution of resistance genotypes separately for the C- and the E-loci.

**Table S19** Resistance allele combinations and their corresponding resistotypes following the genetic model of resistance in *D. magna*. The resistotype describes resistance to *Pasteuria ramosa* C1, C19 and P20 isolates, in that order. The dominant allele at the C-locus confers resistance to C1 and C19, while the dominant allele at the E-locus confers susceptibility to P20. An epistatic relationship links the C- and the E-loci. A "cc" genotype confers susceptibility to P20, regardless of the genotype at the E-locus (see Results section). Highlighted in red is the case where the epistatic relationship is revealed in the phenotype (without it, a "ccee" genotype would underlie a "SS_R" phenotype). Different resistotypes (resistance phenotypes) are grouped with the same background color.

| | CC | Cc | cc |
|---|---|---|---|
| EE | CCEE<br>**RR␣S** | CcEE<br>**RR␣S** | ccEE<br>**SS␣S** |
| Ee | CCEe<br>**RR␣S** | CcEe<br>**RR␣S** | ccEe<br>**SS␣S** |
| ee | CCee<br>**RR␣R** | Ccee<br>**RR␣R** | ccee<br>**SS␣S** |

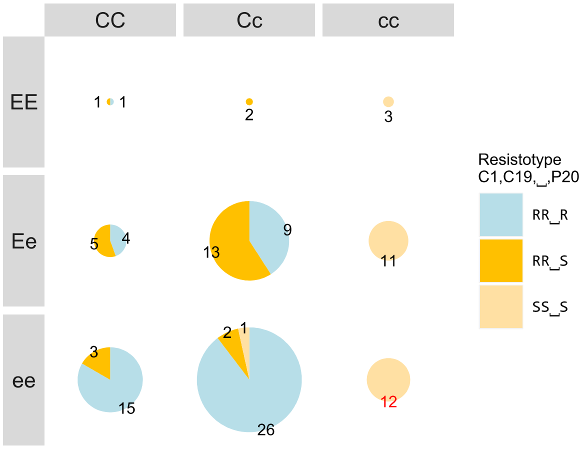

**Figure S4** Distribution of resistance genotypes and phenotypes in the *D. magna* Aegelsee spring population. Data represents individuals from a random sample in spring 2015 (n = 108). Individuals were genotyped using the genetic markers DMPR1 and 3, described in Supplementary GWAS methods, and phenotyped using the attachment test described in the Methods section. Pie charts represent the proportion of resistotypes within each genotype. The size of the charts is proportional to the number of individuals in each genotype group. Mismatches between the expected (Table 1) and observed resistotypes occur when the plots display more than one category. Figure was made using R-packages ggmap (Khale and Wickham 2013) and ggrepel (Slowikowski 2019).

**Table S20** Distribution of genotypes at the C- and the E-loci across resistotypes. Data represents individuals from a random sample in spring 2015 (n = 108). Individuals whose genotypes do not match their resistotype according to the genetic model (see Table 1) are highlighted in grey. The frequency of resistance alleles within resistotypes (highlighted in bold) was used to infer the evolution of resistance allele frequency over the active season (Results section Fig. 6). We calculated allele frequency in the population, and the subsequent expected genotype frequencies at Hardy-Weinberg Equilibrium (HWE). Dominant allele frequency: $p = f(A) = \frac{f(Aa)+2f(AA)}{2n}$; recessive allele frequency: $q = 1 - p$, with genotype frequencies: $f(AA) = p^2$; $f(Aa) = 2pq$ and $f(aa) = q^2$. The frequency of the allele that confers resistance (C- and e-alleles) is displayed.

| | **Cc** | **CC** | **cc** | **C** | **Ee** | **EE** | **ee** | **e** |
|---|---|---|---|---|---|---|---|---|
| **RR␣R** | 35 | 20 | 0 | 75 | 13 | 1 | 41 | 95 |
| ***Frequency*** | *0.636* | *0.364* | *0.000* | ***0.682*** | *0.236* | *0.018* | *0.745* | ***0.864*** |
| **RR␣S** | 17 | 9 | 0 | 35 | 18 | 3 | 5 | 28 |
| ***Frequency*** | *0.654* | *0.346* | *0.000* | ***0.673*** | *0.692* | *0.115* | *0.192* | ***0.538*** |
| **SS␣S** | 1 | 0 | 26 | 1 | 11 | 3 | 13 | 37 |
| ***Frequency*** | *0.037* | *0.000* | *0.963* | ***0.019*** | *0.407* | *0.111* | *0.481* | ***0.685*** |
| **Total** | 53 | 29 | 26 | 108 | 42 | 7 | 59 | 108 |
| **Frequency** | 0.491 | 0.269 | 0.241 | 0.514 | 0.389 | 0.065 | 0.546 | 0.741 |
| **Expected frequency in HWE** | 0.500 | 0.264 | 0.236 | | 0.384 | 0.067 | 0.549 | |
| **Fisher's exact test (using counts)** | | $p = 1$ | | | | $p = 1$ | | |

### Supplementary Selfing methods

#### Selfed *Daphnia magna* parent clones

**Table S21** Overview of the parent clones used to produce the selfed offspring sorted by their inferred genotypes. Resistotype corresponds to the resistance phenotype to the *Pasteuria ramosa* isolates C1, C19 and P20, in that order. The inferred genotypes at the C- and E-loci are the result of the markers analysis of the parent clones (see Supplementary markers Tables S22 to S25). Two exceptions (marked with a "*") are cases where the marker genotype at the E-locus was "EE", although segregation in the selfed F1 offspring revealed a "Ee" genotype. We selfed one to four parents of each homozygous genotype category, and one to four parents of each simple or double heterozygous genotype category. The genotype of the two "SR␣S" parents was inferred using only the segregation pattern of their selfed offspring. The markers were designed to map resistance to C1, C19 and P20 in the Aegelsee population, using the three main resistotypes only: RR␣S, RR␣R and SS␣S, and were consequently not linked to the genotype in SR␣␣ individuals. F0 parents correspond to females caught from plankton (clone name starting with "CH") or hatched from resting eggs (clone name starting with "t") collected from the natural population between 2011 and 2016. Differences in offspring number between the genotype and the phenotype correspond to instances where either genetic markers analysis did not work, or attachment test was not conducted because the *D. magna* clone got extinct.

| Resistotype | Inferred genotype | Repeat name | Parent clone name | n (offspring) Genotype n = 1274 | n (offspring) Phenotype n = 1266 | |
|---|---|---|---|---|---|---|
| RR␣S | CCEE | a | t3_12.3_1i_12 | 43 | 43 | |
| | | b | t3_12.3_1i_21 | 37 | 37 | |
| RR␣S | CCEe | a | CH-2015-36 | 89 | 89 | |
| | | b | t3_12.3_1 | 31 | 31 | |
| | | c | CH-H-2015-49 | 81 | 79 | * |
| RR␣R | CCee | a | CH-H-434-inb2-1 | 39 | 39 | |
| | | b | t1_10.3_2 | 42 | 42 | |
| | | c | CH-H-2015-16 | 66 | 70 | |
| | | d | CH-H-2016-b-70 | 84 | 79 | |
| RR␣S | CcEE | a | t2_17.3_4i_12 | 19 | 19 | |
| RR␣S | CcEe | a | t3_14.3_1 | 47 | 48 | |
| | | b | t2_17.3_4 | 32 | 34 | |
| | | c | t2_17.3_1 | 80 | 64 | |
| | | d | CH-H-2015-59 | 61 | 65 | * |
| RR␣R | Ccee | a | t1_10.3_4 | 36 | 36 | |
| | | b | t5_10.3_3 | 49 | 49 | |
| | | c | CH-H-434-inb2-2 | 21 | 22 | |
| SS␣S | ccEE | a | CH-H-2015-97 | 89 | 87 | |
| SS␣S | ccEe | a | CH-H-2015-113 | 82 | 84 | |
| | | b | t4_10.3_16 | 63 | 65 | |
| SS␣S | ccee | a | t4_13.3_2 | 76 | 74 | |
| | | b | CH-H-2015-86 | 36 | 35 | |
| SR␣S | BbccEE | a | t0_9.3_7 | 35 | 37 | |
| SR␣S | BbccEe | a | t0_28.2_43 | 36 | 38 | |

Supplementary Peas

Genetic model for resistance implemented using the “peas” R-package. The R package “peas” is available at https://github.com/JanEngelstaedter/peas. We present two models, in the first one we consider variation at the C- and E-loci, and in the second one we consider variation at the B-, C- and E-loci.

**Document S1** Implementation of the genetic model of resistance using the “peas” package. Variation at the C- and E-loci. In **step 2.1** we define the genetic system as two loci: C and E presenting each two alleles. Both loci sit on different linkage groups. In **step 2.2** we set the genetic model itself, defining resistance genotypes and their corresponding resistance phenotypes (resistotypes). Resistotypes are presented as resistant (R) or susceptible (S) to the bacteria strains C1, C19 and P20, in that order. Resistance to the bacteria is dominant for the C-locus (Metzger et al. 2016) whereas resistance is recessive at the E-locus. The dominant allele at the C-locus confers resistance to C1 and C19. The recessive allele at the E-locus confers resistance to P20. The epistatic interaction acts between the C- and the E-loci. Recessive individuals at C-locus are susceptible to P20, regardless of their genotype at the E-locus. The genetic model produces 9 possible genotypes coding for 3 different resistotypes. The number of possible genotypes corresponds to the draw of one element among three (AA, Aa or aa genotype) two consecutive times (at two different loci): $n(possible\ genotypes) = 2\binom{3}{1} = 9$. In **step 3** we produce the expected results from the selfing of F0 mothers with all possible combinations of the two loci C and E and store them in a spreadsheet (Results section Table 1 & Supplementary selfing results: Tables S4 to S12).

#### Genetic model for resistance to P20 in the Aegelsee

implemented in peas R package

### 1. Install the package

```
#install.packages("devtools")
#devtools::install_github("JanEngelstaedter/peas", build_vignettes = TRUE)

library(peas)
```

#### C_E genetic model of resistance

##### 2. Set up the genetic model

##### 2.1 Defining the genetic system

```
# 2 loci with 2 alleles each, on different chromosomes
CE <- newGenopheno(nloci = 1,
             alleleNames = list(c("c", "C")))
CE <- addLinkageGroup(CE, alleleNames = list(c("e", "E")))
CE
## Genetic system comprising 2 linkage groups:
##   Linkage group 1: autosomal, 1 locus
##     Alleles at locus 1: c, C
##   Linkage group 2: autosomal, 1 locus
##     Alleles at locus 1: e, E
## No phenotypes defined.
```

##### 2.2 Set genotypes and their corresponding phenotypes: THE GENETIC MODEL

```
CE <- setPhenotypes(CE, "S/R", "__|__","SS_R") # default all recessive --> SS_R
# (we don't take the epistasis relation into account, this will be the last line of the model)

CE <- setPhenotypes(CE, "S/R", "C_|__","RR_R") # C --> R to C1 and C19

CE <- setPhenotypes(CE, "S/R", "C_|E_","RR_S") # E --> S to P20
```

```
CE <- setPhenotypes(CE, "S/R", "cc | __","SS_S") # cc --> S to P20, hides E
```

*Summary of model*

```
CE
## Genetic system comprising 2 linkage groups:
##   Linkage group 1: autosomal, 1 locus
##     Alleles at locus 1: c, C
##   Linkage group 2: autosomal, 1 locus
##     Alleles at locus 1: e, E
## Phenotypes defined for the following traits:
##   S/R (trait values: SS_S, RR_R, RR_S)
```

*List of all possible genotype combinations and corresponding phenotypes*

```
CEallgeno<-getPhenotypes(CE)
CEallgeno # all possible combinations
##         S/R
## cc | ee SS_S
## Cc | ee RR_R
## CC | ee RR_R
## cc | Ee SS_S
## Cc | Ee RR_S
## CC | Ee RR_S
## cc | EE SS_S
## Cc | EE RR_S
## CC | EE RR_S
nrow(CEallgeno) # 9 possible genotypes
## [1] 9
nrow(unique(CEallgeno)) # 3 possible resistotypes
## [1] 3
```

#### 3. Predict crosses

```
CEcombi<-row.names(CEallgeno) # all possible geno combinations from the model
CEcombi<-c(CEcombi[9],CEcombi[6],CEcombi[3],
        CEcombi[8],CEcombi[5],CEcombi[2],
        CEcombi[7],CEcombi[4],CEcombi[1]) # reorder them (optional)
allcrossCE<-list() # create empty list to  store the crossing results

library(xlsx)
# for all the nine possible combinations with C and E varying (F0 mothers)
# we calculate the genotypes and phenotypes fractions in the F1 offspring with the predictCross
function (from the selfing of F0 mothers)

for (i in 1:9) {
# using the "CE" genetic model, we cross the "i" combination with itself
 allcrossCE[[i]]<-predictCross(CE, CEcombi[i], CEcombi[i])

# we add the corresponding resistotypes to the genotype output
 allcrossCE[[i]]$genotypes$SRtrait <- getPhenotypes(CE, equivalent =
"none")[row.names(allcrossCE[[i]]$genotypes), ]

# we store the result in a spreadsheet (genotype output)
 write.xlsx(allcrossCE[[i]]$genotypes,"CEcross.xlsx", sheetName= paste(CEcombi[i],"geno",
as.character(i)), append=T)
```

```
# then the phenotype output in another sheet of the same document
 write.xlsx(allcrossCE[[i]]$phenotypes,"CEcross.xlsx", sheetName= paste(CEcombi[i],"pheno",
as.character(i)), append=T)
 }
```

```
allcrossCE
## [[1]]
## [[1]]$genotypes
##        fraction SRtrait
## CC | EE       1    RR_S
##
## [[1]]$phenotypes
##   S/R fraction
## 1 RR_S       1
##
##
## [[2]]
## [[2]]$genotypes
##        fraction SRtrait
## CC | EE     0.25    RR_S
## CC | Ee     0.50    RR_S
## CC | ee     0.25    RR_R
##
## [[2]]$phenotypes
##   S/R fraction
## 1 RR_R     0.25
## 2 RR_S     0.75
##
##
## [[3]]
## [[3]]$genotypes
##        fraction SRtrait
## CC | ee       1    RR_R
##
## [[3]]$phenotypes
##   S/R fraction
## 1 RR_R       1
##
##
## [[4]]
## [[4]]$genotypes
##        fraction SRtrait
## CC | EE     0.25    RR_S
## Cc | EE     0.50    RR_S
## cc | EE     0.25    SS_S
##
## [[4]]$phenotypes
##   S/R fraction
## 1 RR_S     0.75
## 2 SS_S     0.25
##
##
## [[5]]
## [[5]]$genotypes
##        fraction SRtrait
## CC | EE   0.0625    RR_S
## Cc | EE   0.1250    RR_S
## CC | Ee   0.1250    RR_S
## Cc | Ee   0.2500    RR_S
## cc | EE   0.0625    SS_S
## cc | Ee   0.1250    SS_S
## CC | ee   0.0625    RR_R
## Cc | ee   0.1250    RR_R
## cc | ee   0.0625    SS_S
##
## [[5]]$phenotypes
##   S/R fraction
## 1 RR_R   0.1875
## 2 RR_S   0.5625
## 3 SS_S   0.2500
##
##
## [[6]]
## [[6]]$genotypes
##        fraction SRtrait
## CC | ee     0.25    RR_R
## Cc | ee     0.50    RR_R
## cc | ee     0.25    SS_S
##
## [[6]]$phenotypes
##   S/R fraction
## 1 RR_R     0.75
## 2 SS_S     0.25
##
##
## [[7]]
## [[7]]$genotypes
##        fraction SRtrait
## cc | EE       1    SS_S
##
## [[7]]$phenotypes
##   S/R fraction
## 1 SS_S       1
##
##
## [[8]]
## [[8]]$genotypes
##        fraction SRtrait
## cc | EE     0.25    SS_S
## cc | Ee     0.50    SS_S
## cc | ee     0.25    SS_S
##
## [[8]]$phenotypes
##   S/R fraction
## 1 SS_S       1
##
##
## [[9]]
## [[9]]$genotypes
##        fraction SRtrait
## cc | ee       1    SS_S
##
## [[9]]$phenotypes
##   S/R fraction
## 1 SS_S       1
```

### BC_E genetic model of resistance

**Document S2** Implementation of the genetic model of resistance using the "peas" package. Variation at the B-, C- and E-loci. In **step 2.1** we define the genetic system as three loci: B, C and E presenting each two alleles. The B and C-loci sit on the same linkage group and recombination rate between them is set to r2 = (1-exp(-2*23.1/100))/2 as calculated in (Metzger et al. 2016). The E-locus sits on a different linkage group. In **step 2.2** we set the genetic model itself, defining resistance genotypes and their corresponding resistance phenotypes (resistotypes). Resistotypes are presented as resistant (R) or susceptible (S) to the bacteria strains C1, C19 and P20, in that order. Resistance to the bacteria is dominant for the B and C-loci (Metzger et al. 2016) whereas resistance is recessive at the E-locus. The dominant allele at the B-locus confers resistance to C19. The first epistatic interaction acts between the B and C-loci. The dominant allele at the C-locus confers resistance to C1 and C19, regardless of the genotype at the B-locus. The recessive allele at the E-locus confers resistance to P20. The second epistatic interaction acts between the B/C-loci and the E-locus. Double recessive individuals at the B and C-loci are susceptible to P20, regardless of their genotype at the E-locus. The genetic model produces 27 possible genotypes coding for 5 different resistotypes. The number of possible genotypes corresponds to the draw of one element among three (AA, Aa or aa genotype) three consecutive times (at three different loci): $n(possible\ genotypes) = 3^{\binom{3}{1}} = 27$. In **step 3** we produce the expected results from the selfing of F0 mothers with all possible combinations of the three loci B, C and E and store them in a spreadsheet. From these we extract the results of the F1 selfed offspring groups produced in the present study (Results section Table 2 & Supplementary selfing results Table S13 and S14).

#### 2. Set up the genetic model

##### 2.1 Defining the genetic system

```
library(peas)
r2<-(1-exp(-2*23.1/100))/2 ## recombination rate between B and C loci (Metzger et al. 2016)

# 3 loci with 2 alleles each, BC clustered together (Metzger et al. 2016).
BCE <- newGenopheno(nloci = 2,
             alleleNames = list(c("b", "B"), c("c", "C")),
             rec = r2)
BCE <- addLinkageGroup(BCE, alleleNames = list(c("e", "E")))
```

##### 2.2 Set genotypes and their corresponding phenotypes: THE GENETIC MODEL

```
BCE <- setPhenotypes(BCE, "S/R", "__~__ | __","SS_R") # default all recessive --> SS_R
# (we don't take the epistasis relation into account, this will be the last line of the model)

BCE <- setPhenotypes(BCE, "S/R", "B_~__ | __","SR_R") # B --> R to C19

BCE <- setPhenotypes(BCE, "S/R", "B_~__ | E_","SR_S") # E --> S to P20

BCE <- setPhenotypes(BCE, "S/R", "__~C_ | __","RR_R") # C hides B and --> R to C1 and C19

BCE <- setPhenotypes(BCE, "S/R", "__~C_ | E_","RR_S") # E --> S to P20

BCE <- setPhenotypes(BCE, "S/R", "bb~cc | __","SS_S") # bb and cc --> S to P20, hides E
```

##### *Summary of model*

```
BCE
## Genetic system comprising 2 linkage groups:
##   Linkage group 1: autosomal, 2 loci with recombination rate 0.1849888
##     Alleles at locus 1: b, B
##     Alleles at locus 2: c, C
##   Linkage group 2: autosomal, 1 locus
##     Alleles at locus 1: e, E
## Phenotypes defined for the following traits:
##   S/R (trait values: SS_S, SR_R, RR_R, SR_S, RR_S)
```

*List of all possible genotype combinations and corresponding phenotypes*

```
BCEallgeno<-getPhenotypes(BCE)
nrow(BCEallgeno) # 27 possible genotypes
## [1] 27
nrow(unique(BCEallgeno)) # 5 possible resistotypes
## [1] 5
```

#### 3. Predict crosses

```
BCEcombi<-row.names(BCEallgeno) # all possible geno combinations from the model
allcrossBCE<-list() # create empty list to  store the crossing results

library(xlsx)

for (i in 1:27) {
# using the "BCE" genetic model, we cross the "i" combination with itself
  allcrossBCE[[i]]<-predictCross(BCE, BCEcombi[i], BCEcombi[i])

# we add the corresponding resistotypes to the genotype output
  allcrossBCE[[i]]$genotypes$SRtrait <- getPhenotypes(BCE, equivalent =
"none")[row.names(allcrossBCE[[i]]$genotypes), ]

# we store the result in a spreadsheet (genotype output)
  write.xlsx(allcrossBCE[[i]]$genotypes,"BCEcross.xlsx", sheetName=
paste(BCEcombi[i],"geno", as.character(i)), append=T)

# then the phenotype output in another sheet of the same document
  write.xlsx(allcrossBCE[[i]]$phenotypes,"BCEcross.xlsx", sheetName=
paste(BCEcombi[i],"pheno", as.character(i)), append=T)
}
```

```
# get a specific crossing

match("Bb~cc | Ee", BCEcombi) # position
11
## [1] 11
allcrossBCE[[11]]
## $genotypes
##          fraction SRtrait
## BB~cc | EE   0.0625    SR_S
## Bb~cc | EE   0.1250    SR_S
## BB~cc | Ee   0.1250    SR_S
## Bb~cc | Ee   0.2500    SR_S
## bb~cc | EE   0.0625    SS_S
## bb~cc | Ee   0.1250    SS_S
## BB~cc | ee   0.0625    SR_R
## Bb~cc | ee   0.1250    SR_R
## bb~cc | ee   0.0625    SS_S
##
## $phenotypes
##    S/R fraction
## 1 SR_R   0.1875
## 2 SR_S   0.5625
## 3 SS_S   0.2500
```

```
match("Bb~cc | EE", BCEcombi) # position
20

## [1] 20

allcrossBCE[[20]]

## $genotypes
##          fraction SRtrait
## BB~cc | EE    0.25    SR_S
## Bb~cc | EE    0.50    SR_S
## bb~cc | EE    0.25    SS_S
##
## $phenotypes
##    S/R fraction
## 1 SR_S     0.75
## 2 SS_S     0.25
```

#### Supplementary Markers

Markers developed around GWAS peaks and further used to genotype selfed offspring at the C- and E-loci.

DMPR (*Daphnia magna* - *Pasteuria ramosa*) markers information. Map 2.4 corresponds to the currently available *D. magna* draft genome version 2.4 (Routtu et al. 2014). DMPR1 and 2 are physically linked to the C-locus while DMPR3 and 4 are physically linked to the E-locus. According to the genetic model underlying resistance to *P. ramosa* yielded from the GWAS analysis and previous studies (Metzger et al. 2016), resistance is dominant at the C-locus (resistance to C1 and C19) whereas resistance is recessive at the E-locus (resistance to P20). In addition, an epistatic relationship linking both loci confers susceptibility to P20 to an individual that shows susceptibility to C1 and C19, disregarding the individual genotype at the E-locus. This model is described in Fig. 4. In individuals showing resistance to C1 and C19 (RR␣␣ phenotype, "Cc" or "CC" genotype), DMPR1 and 2 display either a heterozygous pattern with an allele linked to the C-locus dominant allele (called R-allele) and a recessive allele (S-allele) ("Cc" genotype) or a homozygous pattern with one (or two: R1 and R2) dominant allele(s) ("CC" genotype). Note that at markers DMPR1, 3 and 4, two R alleles (called R1 and R2) are found, although R1 and R2 of DMPR4 have identical size. In individuals showing susceptibility to C1 and C19 (SS␣␣ phenotype, "cc" genotype), DMPR1 and 2 display a homozygous pattern with an allele linked to the C-locus recessive allele (called S-allele) ("cc" genotype). In individuals showing resistance to C1 and C19 (without epistatic relationship between C- and E-loci) and susceptibility to P20 (RR␣S phenotype, "Ee" or "EE" genotype at E-locus), DMPR3 and 4 display either a heterozygous pattern with an allele linked to the E-locus dominant allele (called S-allele) and a recessive allele (R-allele) ("Ee" genotype) or a homozygous pattern with an allele linked to the E-locus dominant allele (called S-allele) ("EE" genotype). In individuals showing resistance to C1 and C19 (without epistatic relationship between C- and E-loci) and resistance to P20 (RR␣R phenotype, "ee" genotype at E-locus), DMPR3 and 4 display a homozygous pattern with an allele linked to the E-locus recessive allele (called R-allele) ("ee" genotype). In individuals susceptible to C1 and C19 (SS␣␣ phenotype), DMPR3 and 4 can show one of the three patterns described above (corresponding to "Ee", "EE" or "ee" genotype) because of the epistatic relationship linking the C- and the E-loci.

**Table S22** PCR reaction cycles using DMPR1 to 4.

| Temperature | Time | Cycles |
|---|---|---|
| 95 °C | 15 min | |
| 94 °C | 30 sec | |
| 60 °C | 1 min 30 sec | 30 x |
| 72 °C | 1 min 30 sec | |
| 94 °C | 30 sec | |
| 47 °C | 1 min 30 sec | 10 x |
| 72 °C | 1 min 30 sec | |
| 72 °C | 10 min | |
| 8 °C | ∞ | |

**Table S23** DMPR (*Daphnia magna* - *Pasteuria ramosa*) markers information.

**A:** Description of the positions and sizes of the markers. All primers were used in the same master mix. The size of a marker corresponds to the size of the sequence from the beginning of the F-primer to the end of the R-primer on the reference genome (map 2.4): $size\ marker = \ end\ position - start\ position\ of\ the\ marker$.

**B and C:** Description of primers and motifs. Reference motifs correspond to the motifs that are present in the reference genome (map 2.4). The size of an amplicon is calculated as follows: $size\ amplicon = size\ of\ the\ marker - size\ of\ the\ reference\ motif + size\ of\ the\ motif\ of\ interest$. For example, to obtain the size of the R1-allele of DMPR1 we get: $size\ of\ DMPR_{R1} amplicon = size\ of DMPR1 - size\ of\ DMPR1\ reference\ motif + size\ of\ DMPR1_{R1} motif = 118 - 31 + 2 = 89$.

**D:** Expected marker signatures for all possible genotypes. Here we consider alleles as the observable peaks yielded by the marker analysis, as two copies of the same sequence in a homozygous individual will be observed as a single amplicon of a given size. A “resistance” (R) or “susceptibility” (S) allele does not necessarily confer resistance or susceptibility to the individual that carries it. Resistance phenotype will depend on the genetic model described above. For DMPR1, 3 and 4, we expect R homozygous individuals (“CC” or “ee” genotype) to present one of the two R-alleles (R1 or R2) or both. Heterozygous individuals (“Cc” or “Ee” genotype) should present one R-allele (R1 or R2) together with the S-allele. Finally, S homozygous individuals (“cc” or “EE” genotype) should present only the S-allele. Similarly, for DMPR2, we expect R homozygous individuals to present only the R-allele (R1), heterozygous individuals to present the R-allele (R1) together with the S-allele and S homozygous individuals should present the S-allele. Note that for DMPR4, because both R1- and R2-alleles have the same size, they will appear as a single peak during the marker analysis.

**A**

| Name of marker | Concentration in master mix (µM) | Locus to which marker is physically linked | Scaffold (map 2.4) | Start position of the marker (map 2.4) | End position of the marker (map 2.4) | Size of the marker (bp) (map 2.4) | Position of the motif (map 2.4) |
|---|---|---|---|---|---|---|---|
| DMPR1 | 0.1 | C | 944 | 1350608 | 1350726 | 118 | 1350644 |
| DMPR2 | 0.1 | C | 944 | 1563359 | 1563543 | 184 | 1563391 |
| DMPR3 | 0.6 | E | 2167 | 19662 | 19874 | 212 | 19793 |
| DMPR4 | 0.6 | E | 2560 | 81434 | 81570 | 136 | 81471 |

**B**

| Name of marker | Sequence of forward (F) primer | F-Primer size (bp) | Sequence of reverse (R) primer | R-Primer size (bp) | Melting temperature (°C) | Reference motif (RM) (map 2.4) | RM length (bp) |
|---|---|---|---|---|---|---|---|
| DMPR1 | ACAGCACGTCTCCGACTAAGG | 21 | GACGCCAAMAMCTACGCAACC | 21 | 50 | AAACGCACGGATCCTATATG TATCGAGCTTA | 31 |
| DMPR2 | CAAATCTGCAATGGAATGAAAG | 22 | AACGCAACCGTTACGGTTAC | 20 | 50 | CTCCTGCTGGCT | 12 |
| DMPR3 | TTACGTTCGGTTTGGCTCCG | 20 | TGAAACATTGGTAAGAGACG | 20 | 48 | TACAACAACAACAACAACAA CAA | 23 |
| DMPR4 | GATAGATATTTATTGAACAG | 20 | TTTTGTCTTCGGAAGAACG | 19 | 48 | AATGCCTCCATGCCTCCATGC CTCCA | 26 |

**C**

| Name of marker | Motif of allele(s) conferring resistance (R1 and R2) (dominant at C-locus and recessive at E-locus) | Motif R1 length (bp) | Motif R2 length (bp) | Motif of allele conferring susceptibility (S) (recessive at C-locus and dominant at E-locus) | Motif S length (bp) | Size of R1 amplicon (bp) | Size of R2 amplicon (bp) | Size of S amplicon (bp) |
|---|---|---|---|---|---|---|---|---|
| DMPR1 | AA; AAATGCATATGTATATATCGAGCTTA | 2 | 26 | AAACGCACGGATCCTATATGTATCGAG CTTA | 31 | **89** | **113** | **118** |
| DMPR2 | ATCCTGCTGGCT | 12 | NA | ATCG | 4 | **184** | NA | **176** |
| DMPR3 | TACAACAACAACAACAACAAAA ; TACAACAACAACAACAACAACAAA | 22 | 23 | TACAACAACAACAAAAA | 17 | **211** | **212** | **206** |
| DMPR4 | AATGCCTCCATGCCTCCA ; AATACCTCCATGCCTCCA | 18 | 18 | AATGCCTCCATGCCTCCATGCCTCCA | 26 | **128** | **128** | **136** |

**D**

| Name of marker | Expected marker signature of R homozygous individual ("CC" or "ee" genotype) | Expected marker signature of heterozygous individual ("Cc" or "Ee" genotype) | Expected marker signature of S homozygous individual ("cc" or "EE" genotype) |
|---|---|---|---|
| DMPR1 | R1 / R2 or R1 + R2 | R1 + S or R2 + S | S |
| DMPR2 | R1 | R1 + S | S |
| DMPR3 | R1 / R2 or R1 + R2 | R1 + S or R2 + S | S |
| DMPR4 | R1 / R2 or R1 + R2 | R1 + S or R2 + S | S |
